# Identification of a sub-population of synovial mesenchymal stem cells with enhanced treatment efficacy in a rat model of Osteoarthritis

**DOI:** 10.1101/2024.11.10.622893

**Authors:** Nedaa Al-Jezani, Asmaa Affan, Catherine Leonard, Nabangshu Das, Luiz Gustavo Almeida, Daniel Young, Anand O. Masson, Antoine Dufour, Paul Salo, Pam Railton, James N. Powell, Roman J. Krawetz

**Author notes:** Corresponding Author: Dr. R. Krawetz, McCaig Institute for Bone and Joint Health Faculty of Medicine, University of Calgary, 3330 Hospital Drive NW, Calgary, Alberta, Canada T2N 4N1.

## Abstract

Osteoarthritis (OA) is a painful and debilitating disease which has no cure and there are no treatments which can predictably stop/reverse its progression. Treating this disease is particularly difficult since the articular cartilage lacks intrinsic repair capacity even though mesenchymal stem cells (MSCs) are present in the joint environment and have robust chondrogenic potential. We have previously shown that there is heterogeneity of MSC sub-types within the human synovium, yet it remains unclear if any of these MSC types can regenerate cartilage and/or impact OA disease progression. Therefore, we have undertaken this study focusing on clonally derived MSC populations derived from the synovium of normal and OA patients to characterize if any MSC populations can positively impact OA disease trajectory in a rat model of OA.

MSCs were clonally isolated by indexed flow cytometry, expanded in culture and then characterized for differentiation capacity and by quantitative proteomics. MSC clones were then transplanted into a xenograft rat OA model and treatment effect was determined by histology and immunofluorescence outcomes. We identified heterogeneity in putative MSCs derived from within and between patient groups (normal vs. OA) and the ability of these cells to effect repair in a rat OA model. However, these different sub-types of MSCs could not be distinguished by traditional cell surface markers showing the need for a better understanding of these populations at the single cell level. Using an unbiased proteomics approach, CD47 was identified a novel marker of human MSCs. Using the same rat model of OA, CD47^Hi^ expressing cells were found to have robust treatment efficacy and directly contributed to the formation of new articular cartilage tissue. Characterizing MSCs is essential to understand which sub-types are appropriate for further clinical investigation. If OA patients still have functional MSCs in their synovium, then it is possible these cells can be exploited for cartilage regeneration / OA treatment strategies.

## Introduction

Osteoarthritis (OA) is a chronic joint disease that is characterized by progressive degeneration of the articular cartilage and results in structural changes throughout the joint including: deformity in the subchondral bone, loss of muscle tissue, and inflammation in the synovial membrane(Arden et al., 2020; Grandi and Bhutani, 2020; Ni et al., 2020). These alterations to joint tissues ultimately lead to disability which in turn results in a major socio-economic burden for impacted individuals and their respective healthcare system(Fellows et al., 2016; Palazzo et al., 2014; Salmon et al., 2016). Developing a therapy for OA is particularly challenging for multiple reasons including the multifactorial and elusive nature of the disease(Hafsi et al., 2019; Mora et al., 2018); and inherent properties of the cartilage (absence of blood vessels and nerves) which results in limited endogenous repair capacity(Fellows et al., 2016; Fernandes et al., 2018). All current approved treatments for OA patients focus on symptoms such as pain management, but fail to address the core issue of OA; the degeneration of the cartilage and related structural changes within the joint. In view of these limitations in treatment options, researchers have extensively investigated the use of Mesenchymal stem cells (MSCs) (also referred to as mesenchymal progenitor/stromal cells)(DiMarino et al., 2013) as an alternative treatment modality since they are hypothesized to reduce pain and inflammation through immunomodulatory abilities(Lee et al., 2015), while also potentially being able to induce the repair/regeneration of cartilage through direct differentiation into chondrocytes(Mak et al., 2016) and/or the release of trophic factors to stimulate indirect repair(Soland et al., 2013). MSCs have garnered significant attention over the past two decades; not only because these self-renewing cells possess multi-lineage differentiation capacity but also because these cells have the ability to influence their microenvironment through multiple means(de Witte et al., 2018; Kim et al., 2018).

MSCs have been identified within nearly every mesodermally derived tissue, yet synovial derived-MSCs have been shown to have superior chondrogenic and self-renewing capacity when compared to MSCs derived from a variety of tissues such as bone marrow, and adipose tissue(De Bari et al., 2001; Futami et al., 2012; Nishimura et al., 1999; Sakaguchi et al., 2005). Although MSC-based therapies in numerous pre-clinical studies have shown promising results for the treatment of cartilage injury and/or OA(Jia et al., 2018; Krawetz et al., 2023; Mak et al., 2016; Satué et al., 2019), outcomes from clinical trials have largely been disappointing, with some studies demonstrating that MSC therapy offers little to no advantage over conventional surgical treatments such as microfracture or corticosteroid injection (Koh et al., 2016; Mautner et al., 2023).

In the vast majority of these pre-clinical and clinical studies, cells are isolated and defined as MSCs based on the guidelines proposed by the International Society for Cellular Therapy (ISCT)(Dominici et al., 2006). The minimum criteria that must be met includes adherence to plastic cell culture-ware, multipotent differentiation capacity (typically bone, cartilage and fat), and the expression (and/or lack of expression) of a panel of cell surface markers. Unfortunately, even when satisfying these criteria, MSC phenotypic heterogeneity has been observed in animal and human model systems across multiple tissue types(Phinney, 2012; Russell et al., 2010; Zhou et al., 2019). In a previous study we found significant MSC heterogeneity in clonal populations of cells derived from hip synovium even when cell populations expressed similar combinations of cell surface markers(Affan et al., 2019). Therefore, there is a concern that this heterogeneity might negatively impact the capacity of MSCs to demonstrate clinical efficacy both within cohorts and across studies if we can’t be sure the same cell type (or sub-type) is being isolated and delivered back to patients. Furthermore, since most of the published human MSC data comes from cultured cells, it is also possible that a strong selection bias is induced by the *in vitro* conditions which modifies the properties and populations (sub-types) of the cultured MSCs. Emerging research supports the concept that DPP4⁺ (also marked by PI16 and CD34) synovial cells function as multipotent stromal progenitors. In postnatal mice, DPP4⁺ mesenchymal progenitor cells (MPCs) give rise to Prg4⁺ synovial lining fibroblasts (SLFs) and adipocytes, contributing to normal joint development and synovial homeostasis, with activation and expansion after osteoarthritis (OA) injury. Single-cell transcriptomic atlases further identify PI16⁺ fibroblasts in perivascular niches as latent reservoirs capable of differentiating into specialized synovial lineages following injury. Together, findings across human and mouse OA (Collins et al., 2023; Knights et al., 2022; Li et al., 2024; Peters et al., 2025; Tang et al., 2024) models underscore the progenitor role of these universal fibroblasts and highlight the need for a better understanding of the sub-types of MSCs within the joint environment (by identification of the markers that correlate with functional properties advantageous for therapy) might allow for the isolation of specific populations which are best suited for cartilage regeneration applications.

Therefore, this study employed a unique approach to examine the cell phenotype of clonally derived synovial MSCs obtained directly from synovial tissue (without an initial cell culture step). We have: 1) characterized the cell surface marker expression of clonal populations *in-situ* vs. *in-vitro* to determine the robustness of ISCT recommended markers; 2) correlated the marker expression with multi-potent differentiation capacity; 3) determined if the cell surface marker expression could predict MSC differentiation potential; 4) examined the ability of different clonal populations to treat OA in a preclinical rat model; and 5) characterized the global proteome expression pattern of MSCs isolated from normal vs. OA joints and also MSCs vs. non-MSCs derived sub-populations.

## Methods

### Study Participants

This study protocol was approved by the University of Calgary Human Research Ethics Board (REB15-0005 and REB15-0880). The n=18 (with an additional n=4 for the validation phase) normal control cadaveric donations were harvested within 4 hours of death and were at least 18 years old with no history of arthritis, joint injury or surgery, no prescription anti-inflammatory medications, no co-morbidities. The n=15 (with an additional n=4 for the validation phase) knee OA sample donors were diagnosed by an orthopedic surgeon at the University of Calgary based on clinical symptoms with radiographic evidence in accordance with American College of Rheumatology criteria (**Table S1**). All OA participants provided written consent to participate. All testing was carried out in accordance with the declaration of Helsinki.

### Experimental Outline

The experimental outline of the study is presented in **Figure S1**.

### Cell Sorting and Flow Cytometry

Synovial cells were isolated as previously described(Affan et al., 2019). Briefly, 5 mm² tissue samples were digested using 1 mg/mL Collagenase IV (Thermo Scientific) at 37°C with shaking for 90 minutes. The resulting cell suspension was filtered using a 70 µm filter and centrifuged at 5000 rpm for 6 minutes. The cell pellet was then washed with 1x PBS (Lonza-BioWhittaker) and re-suspended in 100 µL of MesenCult™ Media (StemCell Technologies).

Cells were immunostained for mesenchymal stem cell (MSC) markers, including CD90 (Clone 5E10, PE), CD271 (Clone C40-1457, BV421), CD105 (Clone 266, BV650), CD73 (Clone AD2, APC), and CD44 (Clone G44-26, PE-Cy7) (all from BD Bioscience), according to the manufacturers’ protocols. Additionally, cells were stained with the viability marker FVS510 (BV510) at a 1:100 dilution for 30 minutes on ice and with CD68 (Clone Y1/82A, FITC) to target the macrophage population. Positive compensation controls were conducted using Ultracomp eBeads (eBioscience), and unstained cells served as negative controls.

The labeled cells were sorted using a BD FACS Aria Fusion with an indexed sorting protocol. The instrument was configured with a 100 µm sorting nozzle, a “single cell” mask, a 2X neutral density filter, and a flow rate below 50% of the maximum. Dead cells (FVS510^+^) and macrophages (CD68^+^) were excluded, and the remaining cells were collected into 96-well plates (1 cell per well) containing Dulbecco’s modified Eagle’s medium F-12 (DMEM/F-12) with MesenCult™-SF attachment substrate (StemCell Technologies) and 1% antibiotic-antimycotic (Life Technologies). The sorted cells were cultured until they reached 70-80% confluency, then transferred to T-25 flasks (Greiner). Once the cultures reached ∼70% confluency, the cells were washed with 1x PBS (Lonza-BioWhittaker), trypsinized (Corning), and re-plated. Media was changed every 2 days, and cells were passaged until they expanded into 4x T-75 flasks.

During the validation phase of the study, bulk sorting was performed directly from dissociated synovium without prior cell expansion. Cell suspensions from a new cohort (n=4 normal and n=4 osteoarthritis synovium) were isolated using the same digestion method described earlier. The cells were stained with the markers CD68, CD90, CD73, CD44, and the viability marker FVS510, as previously detailed. Sorting was performed using the BD FACS Aria Fusion, and cells were sorted into 5 mL tubes containing 500 µL of media. Instead of sorting one cell per well, the strategy involved gating out CD68+ and dead cells, followed by sorting CD90^+^CD73^+^CD44^+^ cells into one tube and the remaining cells (a mixture of all populations except CD90^+^CD73^+^CD44^+^) into a separate tube. This sorting was conducted using a 100 µm sorting nozzle, a “purity” mask, a 2X neutral density filter, and a flow rate of 5/11. Both positive and negative populations were transferred to 48-well plates and expanded until they reached 70% confluency in 4x T-75 flasks. These cells were subsequently characterized for their differentiation potential and underwent *in vitro* flow cytometry analysis as previously described.

### Differentiation Analysis

Expanded clonal cells were induced to differentiate into bone, cartilage, and fat as previous(Affan et al., 2019; Krawetz et al., 2022). For each differentiation protocol, 5×10⁵ cells were either plated (for osteogenic and adipogenic protocols) or pelleted and plated (for chondrogenic protocols) in 24-well plates. Cells were incubated for 21 days in either osteogenic, chondrogenic, or adipogenic media, with the media changed every 2 days: **Osteogenic media**: DMEM/F-12, 10% FBS, 1% antibiotic-antimycotic, 1% MEM non-essential amino acids (NEAA), 10⁻⁴M dexamethasone, 50µg/mL L-ascorbic acid (AA), and 10mM β-glycerophosphate (all from Sigma). **Chondrogenic media**: DMEM/F-12, 10% FBS, 1% antibiotic-antimycotic, 1% MEM NEAA (Gibco), 10nM dexamethasone, 50mg/mL AA, NaOH, 10ng/mL TGF-β3 (Peprotech), 500ng/mL BMP-2 (Peprotech), sodium pyruvate (Gibco), and insulin-transferrin-selenium (ITS, Lonza-BioWhittaker). **Adipogenic media**: DMEM/F-12, 10% FBS, 1% antibiotic-antimycotic, 1% MEM NEAA, 1μM dexamethasone, 10µM insulin, 200µM indomethacin, and 500µM isobutylmethylxanthine (all from Sigma).

### Quantitative PCR (qPCR)

Cells were washed with 1X PBS. RNA was extracted using TRIzol reagent (Invitrogen) from osteogenic and adipogenic cells, and using the Total RNA Kit I (OMEGA bio-tek) from chondrogenic cells, following manufacturers’ protocols. The isolated mRNA was stored at –80°C for later analysis. To generate cDNA, 10μL of mRNA was mixed with 10μL of cDNA Master Mix (High-Capacity cDNA kit, Applied Biosystems) and incubated in a thermocycler (Bio-Rad) for 2 hours. The samples were then stored at –20°C. qPCR was used to quantify the gene expression of specific markers: **Osteoblast markers**: *Sp7, Runx2*, **Chondrocyte markers**: *Sox9, Col2a*, **Adipocyte markers**: *Adipoq*. Triplicate reactions per sample were performed in either 384-well or 96-well plates (Applied Biosystems). Reactions included 0.5µL of the specific probe, 5µL of TaqMan Universal PCR Master Mix (Applied Biosystems), 3.5µL ultrapure H₂O, and 1µL cDNA. Gene expression was standardized using the 18S ribosomal RNA, calculated via the DeltaDeltaCT method.

### In Vitro Histology

Cells were fixed with 10% neutral buffered formalin (NBF) at room temperature for 1 hour. **Osteogenic cultures** were stained with 20mg/mL alizarin red (Sigma) for 45 minutes. **Adipogenic cultures** were washed, incubated with 60% isopropanol, and stained with Oil Red O for 15 minutes, followed by Harris Hematoxylin. **Chondrogenic cultures** were treated with 1% acetic acid and stained with 0.1% Safranin O for 5 minutes as previously described(Affan et al., 2019).

### Destabilization of the Medial Meniscus (DMM) Injury Model

Ten-week-old Lewis rats were used for the DMM model. Males weighed ∼250g (±50g), and females weighed ∼175g (±20g) at the time of surgery. Rats were anesthetized with isoflurane, and a medial para-patellar arthrotomy was performed under a microscope. The fat pad over the cranial horn of the medial meniscus was retracted, and sectioning of the medial meniscotibial ligament destabilized the meniscus(Das et al., 2023; Iqbal et al., 2016). Sham-operated rats served as controls, receiving surgery without injury. The joint capsule and skin were closed with sutures and adhesive.

### Cell Injection

Once a sufficient number of cells had been obtained, the purified synovial cells were incubated with the following mix overnight (∼12 h) at 37 °C, 2% O_2_: 5 mL MesenCult™, 5 μL tdTomato lentivirus (where tdTomato was driven by the EF1-alpha promoter), and 2 μL of Polybrene (8 μg/mL, Sigma). Medium was changed the following day. The cells were then sorted for tdTomato expression and the positive cells were used for the xenotransplantation experiments.

Control groups included 6 rats with sham surgery and 6 rats that underwent DMM injected with PBS alone. The injection involved a 30G needle placed through the patella tendon into the joint space, with 10µL of sterile DPBS containing 100,000 cells injected 1 week post-DMM surgery. Sample size as follows for the initial experiments: DMM (MSC injected) n=3 rats per cell line, 8 cell lines used (normal MSC x2 and non-MSC x2, OA MSC x2 and non-MSC x2). Sample size as follows for the CD47 experiments: DMM (CD47^Hi^ injected) n=2 rats per cell line, 6 cell lines used (3 normal, 3 OA), (CD47^Lo^ injected) n=2 rats per cell line, 6 cell lines used (3 normal, 3 OA).

### Histology and OA Grading

Four weeks post-cell injection (5 weeks post-DMM), rats were sacrificed. Knee joints were dissected, fixed in 4% formalin, decalcified, embedded in paraffin, and sectioned. Sections were stained with Safranin O to visualize proteoglycans and graded according to OARSI guidelines for rat knees.

Immunohistochemistry analysis was performed on the rat knee sections. Antigen-retrieval was achieved using 10 mM sodium citrate (pH 6.0), and non-specific blocking was prevented using goat-serum (1:500 dilution in TBST). tdTomato (AB8181, Origene), Collagen 2 (Col2; Clone # II-II6B3, DSHB), PRG4 (Clone # 9G3, Millipore), Ccl2 (Clone # 2D8, ThermoFisher), Ki67 (Clone # SolA15, ThermoFisher), Sox9 (Clone # 7H13L8, ThermoFisher) or CD47 (Clone # B6H12, ThermoFisher) were applied to the sections and incubated overnight. For the Col2 staining an additional hyaluronidase (Sigma) treatment step was added. Secondary controls were also performed, where only secondary antibody was applied to the sections (no primary antibody). All slides were mounted using EverBrite™ Hardset Mounting Medium with 4′,6-diamidino-2-phenylindole (DAPI, Biotium). Slides were imaged using a Plan-Apochromat objective on an Axio Scan.Z1 Slide Scanner microscope (Carl Zeiss).

### Tissue Cytometry

TissueQuest software was used for immunofluorescent image analysis. After nuclear segmentation using DAPI, each marker channel was analyzed, with gates and thresholds set using non-stained controls. Data were visualized using scattergrams and histograms, and statistical analysis was done using GraphPad.

### Quantitative Shotgun Proteomics Using Tandem Mass Tags (TMT-6) Labeling

Synovial membrane-derived clones were lysed and sonicated. Cellular proteins were quantified and isolated for TMT-6 plex shotgun proteomics. LC-MS/MS analysis was conducted at the Southern Alberta Mass Spectrometry core facility(Das et al., 2023). Peptides were separated using C18 columns, and data were analyzed using MaxQuant software and statistical tools in R.

### Reactome Pathway Analysis

The STRING database was used to identify protein interconnectivity, with analysis done in Metascape for functional enrichment, interactome analysis, and gene annotation. Human protein data were analyzed with a 1% false discovery rate.

### Statistics

Statistical analysis was conducted using GraphPad Prism 7. Data were presented as mean ± standard deviation (SD). Student’s t-test was used for group comparisons, and Spearman correlation was applied for association analyses. Significance was set at p<0.05, with data visualized using IBM SPSS.

## Results

### Clonal MSC Derivation

In this study, synovial biopsies were collected from 15 patients undergoing knee orthopedic procedures and 18 normal cadaveric donors (**Table S1**). From the normal synovium, 228 clonal cell lines were generated, while 259 clonal lines were derived from osteoarthritic (OA) synovium (**Table S2**). Out of these, 29 clones from the normal synovium and 37 from the OA synovium exhibited the self-renewal potential required to expand to a population size suitable for further analysis, such as differentiation and flow cytometry (**Table S3**).

To complete all differentiation and flow cytometry analyses (including replicates), each clonally derived synovial MSC line needed to undergo approximately 19 population doublings. However, most clonal cell lines lacked the self-renewal capacity necessary to meet the study’s characterization criteria (**Table S3**).

A negative correlation was observed between donor age and the population doublings of clones derived from that donor (**Figure S2**). This relationship was more pronounced in OA patients compared to normal donors. No correlation was found between sex and population doublings (**Figure S2**).

### Cell Surface Marker Expression In Situ vs. In vitro

The data presented is a representative example from one normal and one OA patient (**Figure S3**), while the collective results are summarized in **Tables S4 and S5**. Cell surface marker profiles were compared *in situ* (pre-culture, indexed cell sorting was used to record cell surface profiles) and *in vitro* (post-culture). Flow cytometry analysis revealed that cell surface expression of the cloned cells (isolated from both OA and Normal) became altered by exposure to the cultural microenvironment (**Figure S3 A,B**). Specially, 4 clones were obtained from a single normal individual. Clone 1 was initially positive for CD90, CD44, and CD73 *in situ* and negative for CD105 and CD271, yet this clone gained the expression of CD105 *in vitro*. Clones 2,3 and 4 shared similar expression *in situ* (expressing CD44, and CD73 while lacking the expression of CD90, CD105 and CD271). Similar to clone 1, these clones acquired the expression of CD105 as well as CD90 (**Figure S3 A**). Clones attained from a single OA individual show a similar pattern. Clones 1 and 2 were positive for CD44 and negative for CD90, CD73, CD105 and CD271 *in situ* and gained the expression of CD90, CD73 and CD105 *in vitro*. Clone 3 was CD44 and CD73 positive *in situ* and gained expression of CD90 and CD105 *in vitro*. Clone 4 expressed CD90 and CD73 in situ and gained expression of CD44 and CD105 *in vitro*.

Collectively, the common trend observed in the clones presented and those from the entire cohort was that despite a heterogeneous cell surface marker expression profile *in situ*; the marker profile aligned *in vitro* with nearly all clones (normal and OA) expressing CD90, CD44, CD73, and CD105 (**Figure S3, Tables S4 and S5**). Since this experiment was performed clonally, we can conclusively state that the individual cells self-regulated their unique cell surface profiles to a common profile once expanded in culture. Overall, this demonstrates a convergent selection bias towards the traditional cell surface marker profile expressed by MSCs.

### Multipotential Differentiation Assessment

Aside from cell surface marker expression a hallmark of ‘stemness’ includes self-renewal ability and multipotent differentiation capacity; therefore, clones were identified as MSCs if they demonstrated differentiation into bone, cartilage and fat *in vitro*. The data presented is a representative example from <u>one normal</u> and <u>one OA</u> patient (**Figure 1**), while the collective results summarized in **Tables S4 and S5**. Based on histological analysis, normal clone 1 and 2 stained positive for Alizarin Red, Safranin O, and Oil Red O when induced to differentiate into osteoblasts, chondrocytes and adipocytes (**Figure 1 A**). When qPCR analysis was performed, clone 1 expressed *Sp7* (osteogenic), *Col2A1* and *Sox9* (chondrogenic), but lacked the expression of *Adiponectin* (adipogenic) (**Figure 1 B**). Clone 2 upregulated all differentiation markers (**Figure 1 B**). Clone 3 and 4 lacked Alizarin Red staining but stained positive for both Safranin O and Oil Red (**Figure 1 A**). Clone 3 expressed *Sp7*, *Col2a* and *Adiponectin* (**Figure 3 1**) while clone 4 only expressed *Adiponectin* (**Figure 3 1**).

**Figure 1.**
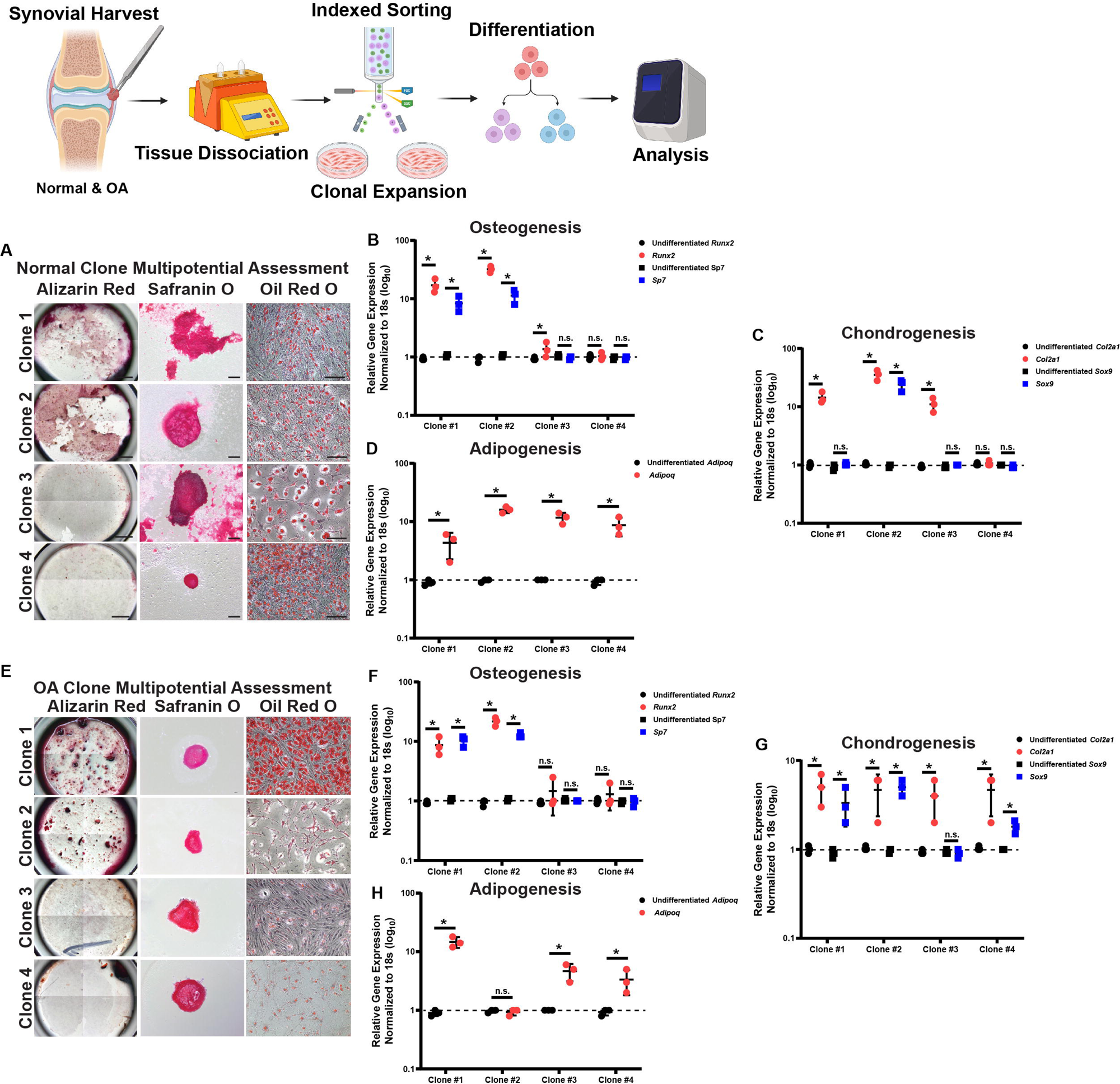
*In vitro* differentiation potential of clones derived from normal individuals and patients with OA. Representative histological staining of Alizarin Red, Safranin O and Oil Red O from 4 normal (A) and 4 OA clones (E). QPCR data (n=3 biological replicates and n=3 technical replicates, each point is the mean of the technical replicates) is presented for each of the clones for osteogenic (B,F), chondrogenic (C,G) and adipogenic (D,H) markers. *p<0.05, **p<0.01

Similar patterns with observed in the clones derived from OA patients. Clone 1, when induced to osteogenesis, chondrogenesis, and adipogenesis, was positive for Alizarin Red, Safranin O and Oil Red (**Figure 1 C**) and expressed *Sp7*, *Runx2*, *Col2a1* and *Adiponectin* (**Figure 1 D**). Clone 2 stained positive for Alizarin Red, Safranin O, Oil Red O and expressed osteogenic and chondrogenic markers but lacked the expression of *Adiponectin* marker (**Figure 1 D**). Clones 3 and 4 were negative for Alizarin Red but stained positive for Safranin O and Oil Red O (**Figure 1 C**), but both clones expressed osteogenic markers (**Figure 1 D**). Clone 3 lacked the chondrogenic markers, clone 4 displayed up-regulation of Sox9 (**Figure 1 D**). Clone 3 expressed *Adiponectin* while clone 4 did not (**Figure 1 D**).

Based on ISCT criteria, for a clone to be considered as an MSC it needed to demonstrate multipotent differentiation capacity in osteoblasts, chondrocytes and adipocytes (Dominici et al., 2006). Since we observed some level of disagreement between histological and molecular analysis of differentiation, we decided that each clone had to demonstrate positive histological staining with positive marker gene expression to be considered conclusively differentiated into a specific lineage Therefore, for the 8 clones presented in **Figure 1**, only normal clones 1, 2 and OA clones 1, 2 were defined as MSCs. **Tables S4 and S5** summarizes all data collected from both normal and OA knee joints respectively. When the differentiation capacity of the clones was compared to the cells *in situ* (pre-culture) cell surface marker expression, the most frequent marker phenotype of cells that met the minimal criteria for MSCs (in situ) was CD90^+^CD44^+^CD73^+^.

### Cell Surface Profile Validation

To determine if CD90^+^CD44^+^CD73^+^ expression defines MSC populations *in situ*, synovial membrane samples from a new cohort of patients (n=4 normal, n=4 OA) were obtained. In this experiment we directly compared between cells that were triple positive for CD90^+^CD44^+^CD73^+^ (live, CD68^-^) vs. all live CD68^-^ cells that did not have this exact expression profile. Interestingly, cells enriched for CD90^+^CD44^+^CD73^+^ expression did not guarantee multipotent differentiation potential (summarized in **Table S6**). On the contrary, samples that did not contain the CD90^+^CD44^+^CD73^+^ population still retained differentiation potential in OA and normal synovium meeting the MSC criteria. Furthermore, after culture *in vitro*, all lines expressed all MSC markers except CD271. Based on this experiment, an expression profile of CD90^+^CD44^+^CD73^+^ on cells directly isolated from synovial membrane does not presumptively identify MSCs.

### In vivo Functional Assessment

It was next decided to investigate if there was a functional difference between MSC vs. non-MSC clonally derived populations when injected into the knees of rats with surgically induced OA. DMM surgery was performed on the left knee of Lewis rats and one-week post DMM surgery, rats were either injected with clones classified as MSCs or non-MSCs (from both normal and OA synovium). Histological sections from control (sham injured) and injured (treated with saline) rats were compared to rats that had been injured and injected with either MSC or non-MSC clones (**Figure 2**). Knees that received sham DMM appeared to be morphologically normal (**Figure 2 A**), while knees that underwent DMM and received saline showed proteoglycan loss, areas of cartilage loss and osteophyte formation (**Figure 2 B**). DMM knees injected with MSCs showed some areas of reduced proteoglycan staining and surface fibrillation but did not present with areas of cartilage loss (**Figure 2 C**). Injured knees that received non-MSCs showed proteoglycan loss, areas of synovial inflammation, surface fibrillation and areas of cartilage loss (**Figure 2 D**). Osteoarthritis Research Society International (OARSI) histological scoring and Krenn scoring were undertaken to quantify OA severity and synovial inflammation respectively. These scores demonstrated that rats receiving MSCs had a significantly lower OARSI and Krenn score vs. rats injected with non-MSCs post-DMM surgery (**Figure 2 E, Figure S4**). These finding suggested that intra-articular injection of MSCs either inhibited degeneration and/or promoted regeneration to reduce the severity of OA, while the non-MSCs lacked this functional ability.

**Figure 2.**
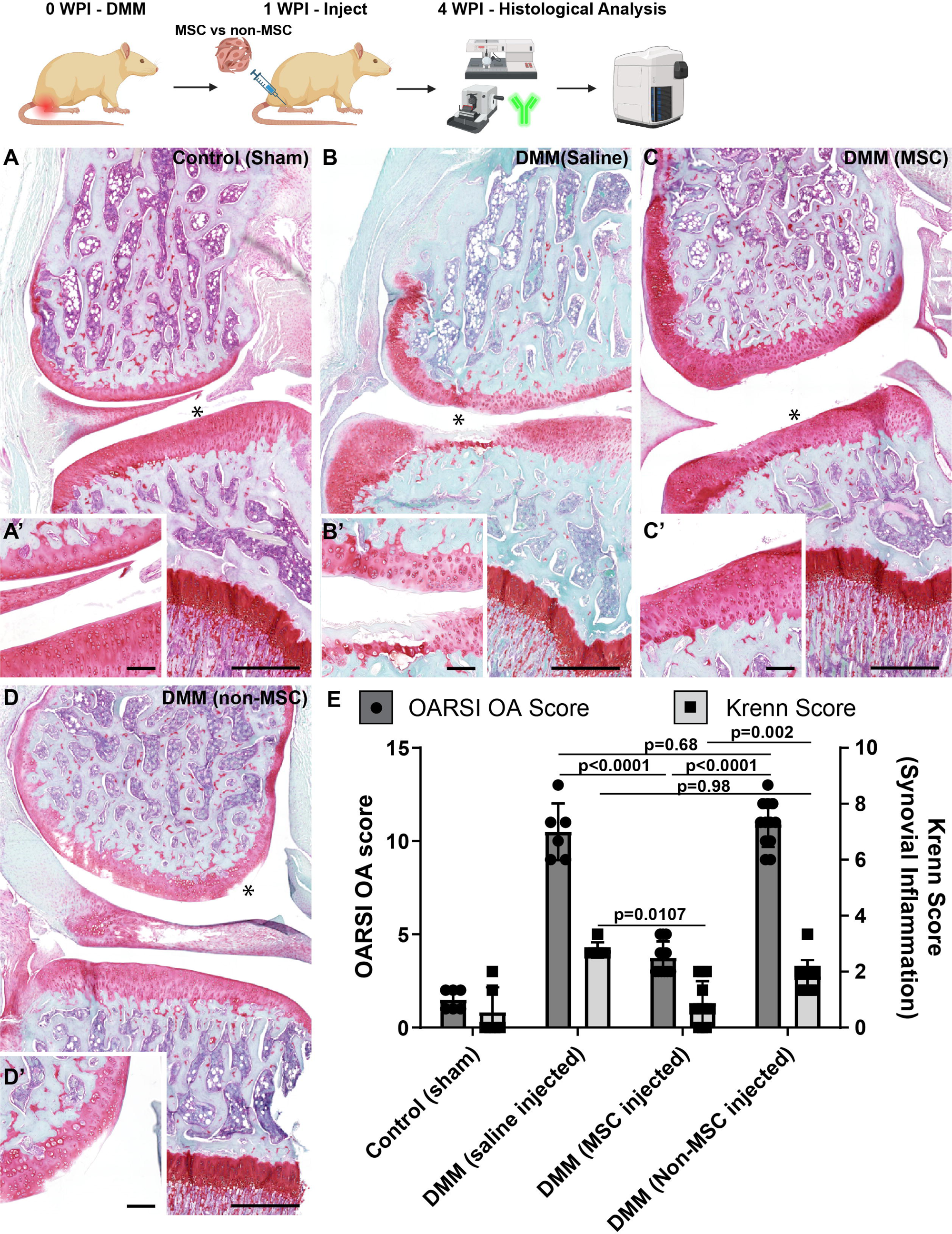
OA scoring with and without cell injection in a rat DMM model. Representative images of rats with sham surgery (A, A’), DMM injury with injection of saline (B, B’), DMM injury with injection of MSCs (C, C’) or non-MSCs (D, D’). The OARSI and Krenn scoring was quantified in each group (E). Scale bars equal 200µm in A,B,C,D and 20µm in A’,B’C’D’. *p<0.05. Sample size as follows: control (sham) n=6, DMM (saline injected) n=6, DMM (MSC injected) n=3 rats per cell line, 8 cell lines used (normal MSC x2 and non-MSC x2, OA MSC x2 and non-MSC x2). Normal and OA MSC groups and non-MSC groups added together.

To examine whether the transplanted cells populations were capable of directly contributing to articular cartilage regeneration *in vivo*, the clones were virally labeled with TdTomato (to track their localization within the rat tissue). TdTomato^+^ MSC clones (from normal and OA patients) were observed within the articular cartilage (**Figure 3 G,H; M,N**) while TdTomato^+^ non-MSC clones (from normal and OA patients) were not observed with cartilage tissue (**Figure 3 J,K; P,Q**). In uninjured rats (**Figure 3 A,B**) or injured rats injected with saline alone (**Figure 3 A,B**), no TdTomato signal was detected.

To determine if the transplanted human cells generated functional articular chondrocytes, Lubricin/PRG4 straining was undertaken, since this protein is produced by chondrocytes and is essential to joint homeostasis in part by lubricating the articular surfaces(Das et al., 2019; Iqbal et al., 2016). In control (sham) joints, lubricin staining is primarily confined to the superficial zone of the cartilage resulting in a continuous layer of expression (**Figure 3 A,C**). In DMM joints injected with saline, this protective layer of PRG4 staining is disrupted (**Figure 3 D,F**). When MSCs (from normal or OA synovium) were injected into DMM joints, some level of restoration of the PRG4 protective layer was observed (**Figure 3 G,I,M,O**). Interestingly, it was observed that normal MSC clones throughout the cartilage produced PRG4 (**Figure 3 G,I**), while this was not observed with OA MSC clones (**Figure 3 M,O**). In DMM joints injected with non-MSC clones (normal of OA), the PRG4 protective layer was disrupted (**Figure 3 J,K,P,R**) similar to saline injected joints (**Figure 3 D,F**). A tissue cytometry approach (**Figure S5**) was used to quantify the number of tdTomato, PRG4 and tdTomato with PRG4 expressing cells in the articular cartilage (**Figure 3 S**). Overall, there was a significant increase in tdTomato and PRG4 expressing cells within the cartilage post-MSC treatment (regardless of being derived from normal/OA synovium); yet more tdTomato and PRG4 expressing cells were observed in joints treated with MSCs derived from normal vs. OA synovium. Furthermore, we also observed that while transplanted normal MSCs took on a PRG4 positive phenotype, MSCs from OA synovium did not. Non-MSCs, regardless of patient population did not engraft into the articular cartilage nor take on a PRG4 positive phenotype (**Figure 3 S**).

To determine if the MSC clones were regenerating/maintaining the articular cartilage in DMM joints, collagen 2 staining was undertaken. Collagen 2 staining was found within the articular cartilage of DMM joints and colocalized with TdTomato expression (**Figure S6**). Taken together, these outcomes suggest that intra-articular injection of MSCs (normal or OA) resulted in the regeneration of articular cartilage in comparison with the intra-articular injection of non-MSCs that had no beneficial effect on the articular cartilage.

### Localization of non-MSC Clones Within the Joint

Since non-MSC clones were not observed within the articular cartilage of the rats, the synovial tissue was examined (**Figure 4**). Few MSCs (normal and OA – normal clone shown as a representative example), were found within the synovium or adjacent tissue (**Figure 4 A**); while many non-MSCs (normal and OA – normal clone – same patient as MSC clone, shown as a representative example) where observed throughout the synovium in both the intimal and subintimal layers(**Figure 4 B**). To determine what role the cells were playing in the micro-environment, CCL2 staining was performed. CCL2 is a pro-inflammatory chemokine which recruits circulating monocytes from the blood to infiltrate tissue during inflammatory processes(Deshmane et al., 2009); is involved in OA pain and pathogenesis(Jablonski et al., 2019; Miller et al., 2012; Raghu et al., 2017). We have demonstrated that CCL2 is expressed in OA MSCs and regulates their behaviour(Harris et al., 2013). Although CCL2 staining did co-localize with tdTomato (MSCs), it was minimal (**Figure 4 A**). CCL2 staining was also observed in the intimal layer but was independent of tdTomato expression (**Figure 4 A**). This was in contrast to CCL2 staining when non-MSC were injected into joints (**Figure 4 B**). In non-MSC injected joints, CCL2 staining was observed throughout the synovium and was highly co-localized with TdTomato expression (**Figure 4 B**). Furthermore, since CCL2 is a secreted protein, it was interesting to observe that CCL2 staining appeared to diffuse out from the tdTomato^+^ non-MSC clones (**Figure 4 B**). When the number of CCL2 positive cells and the level of CCL2 staining (mean fluorescent intensity – MFI) was quantified, it was found that significantly more CCL2 positive cells and expression was detected in the synovium of rats injected with non-MSCs (normal or OA) (**Figure 4 C**).

**Figure 3.**
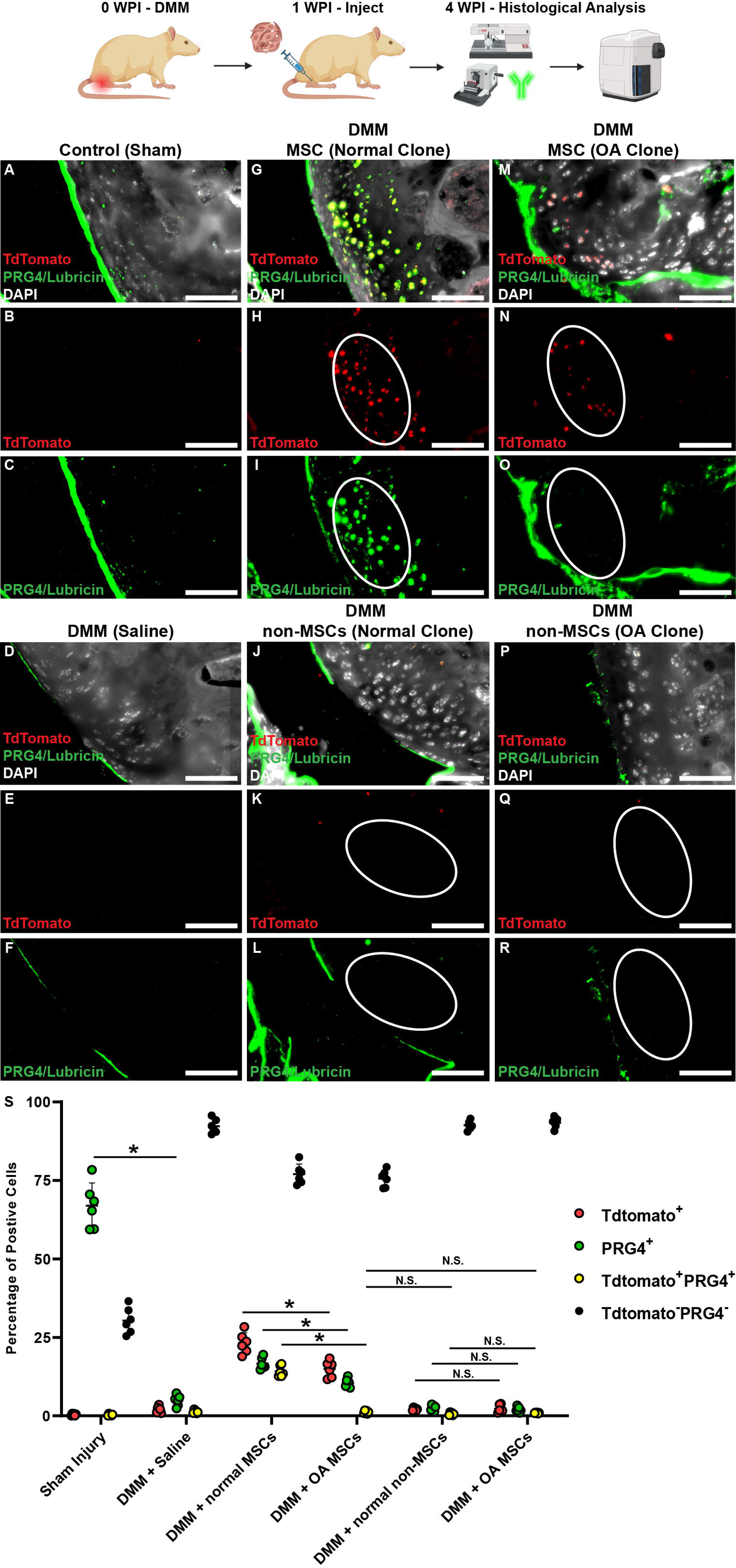
Exogenous cell contribution to cartilage repair in a rat DMM model. Representative images of rats with sham surgery (A) without cell injection (no TdTomato – B) and endogenous PRG4/Lubricin staining (C). In rats injected with saline (D), no TdTomato (E) and disrupted PRG4/Lubricin (F) staining was observed. When normal (G) or OA (M) MSCs where injected into injured rats, TdTomato expression (H.N) was observed. When normal (J) or OA (P) non-MSCs were injected no TdTomato (K,Q) and disrupted PRG4/Lubricin (L,R) staining was observed. Scale bars equal 30µm. Tissue cytometry was employed to quantify the number of tdTomato positive, PRG4 positive and double positive cells within the articular cartilage (S). *=p<0.05, N.S. = not significant. Sample size as follows: control (sham) n=6, DMM (saline injected) n=6, DMM (MSC injected) n=3 rats per cell line, 8 cell lines used (normal MSC x2 and non-MSC x2, OA MSC x2 and non-MSC x2).

**Figure 4.**
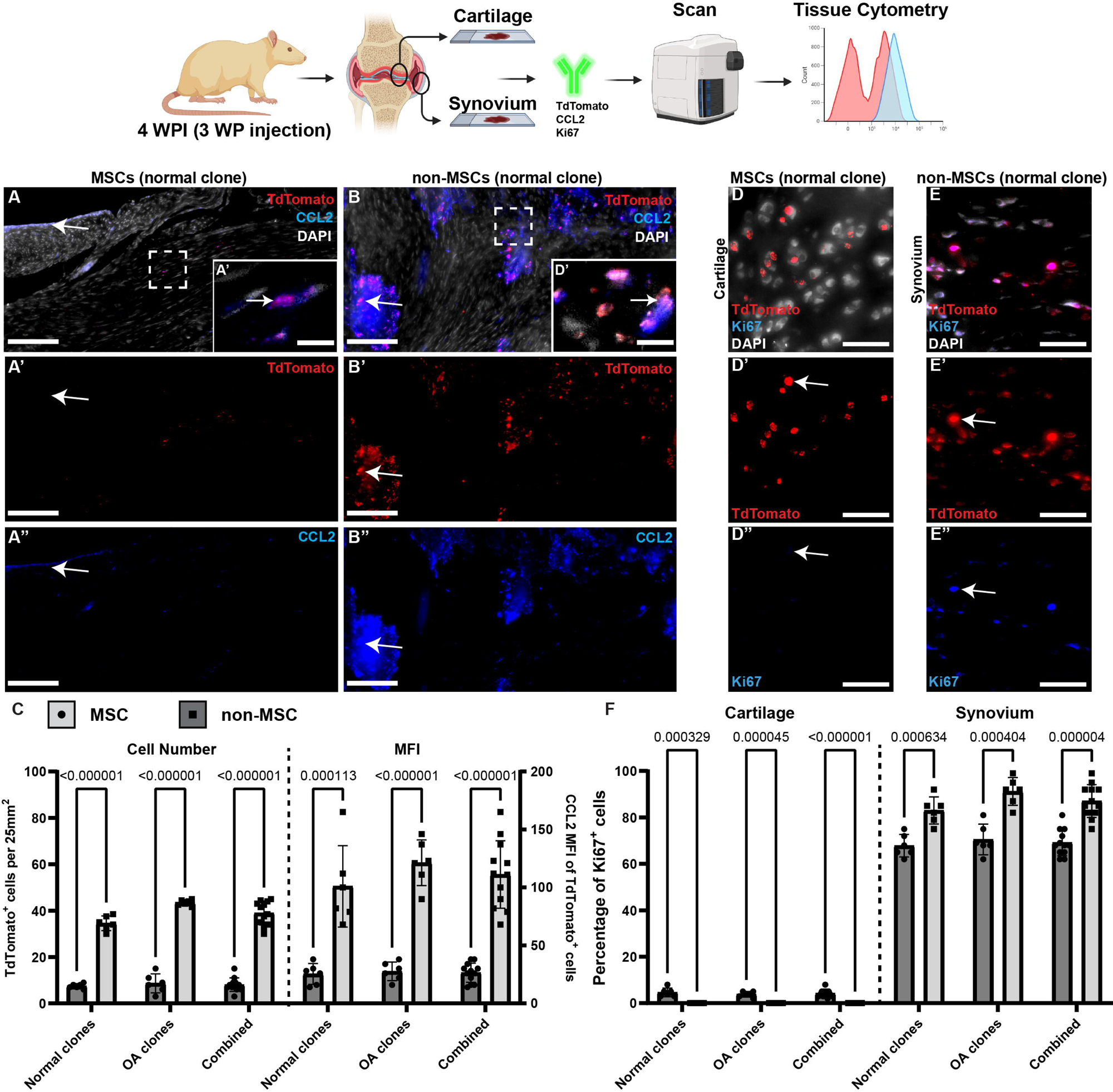
Contribution of transplanted cells to inflammatory signaling. Representative images of MSCs (A) and non-MSCs (B) in the synovium of rats post-DMM. In rats injected MSCs, only minimal CCL2 (A”) expression is observed and/or colocalized with tdTomato staining (A’). In rats injected with non-MSCs, intense CCL2 staining (B”) was observed that colocalized with tdTomato staining (B’). this was quantified using tissue cytometry (C). In rats injected MSCs, little Ki67 staining was observed in the cartilage (D-D”). In rats injected with non-MSCs, nearly every transplanted cell in the synovium was also positive for Ki67 (E-E”). This was quantified using tissue cytometry (F). *=p<0.05, n.d. = none detected.

In a previous study, we demonstrated that CCL2 acted upon MSCs by increasing their proliferative ability(Harris et al., 2013). Therefore, we decided to examine MSC/non-MSC proliferation in the joint (synovium and cartilage) through Ki67 staining (**Figure 4 D-F**). Within the articular cartilage, transplanted MSCs (normal or OA) demonstrated little to no Ki67 staining indicating minimal cell proliferation (**Figure 4 D**). Since non-MSC clones were not observed in the articular cartilage, no Ki67 staining was observed. In the synovium, MSC clones (normal or OA) demonstrated minimal positive Ki67 staining, while non-MSC clones demonstrated robust staining (**Figure 4 D**). However, it is important to remember that significantly fewer MSCs were observed within the synovial tissue (**Figure 4 G**), yet these cells were proliferative.

### Proteomics Analysis of clonal populations

Since a clear difference was observed between *in vitro* and *in vivo* functionality/behaviour in the two different populations of synovial cells; a quantitative shotgun proteomics analysis was undertaken to understand the proteomic differences between the sub-types (**Figure 5A**). Several differentially expressed proteins were identified between the sub-types (**Table S7, Figure 5B**). Using Metascape (**Figure 5C**) and STRING-db (**Figure 5D**) analyses were undertaken on the proteins differentially regulated in MSCs, several pathways were differentially enriched. An aspect of this proteomic workflow that was interesting was that the MSC sub-type was enriched for CD47, ITGA5 and DPP4 expression. While ITGA5 and DPP4 have already been implicated in fibroblast and progenitor function, CD47 has been traditionally thought of as a signal to immune/macrophages that blocks their phagocytosis of these CD47 expressing cells. In terms of MSC expression of CD47, this molecule has been shown to enhance homing and immunomodulatory ability of the cells. Therefore, we characterized freshly isolated synovial samples from normal (n=3) and OA (n=3) joints for CD90^+^CD73^+^CD44^+^ cells and examined what percentage of this population expressed CD47.

**Figure 5.**
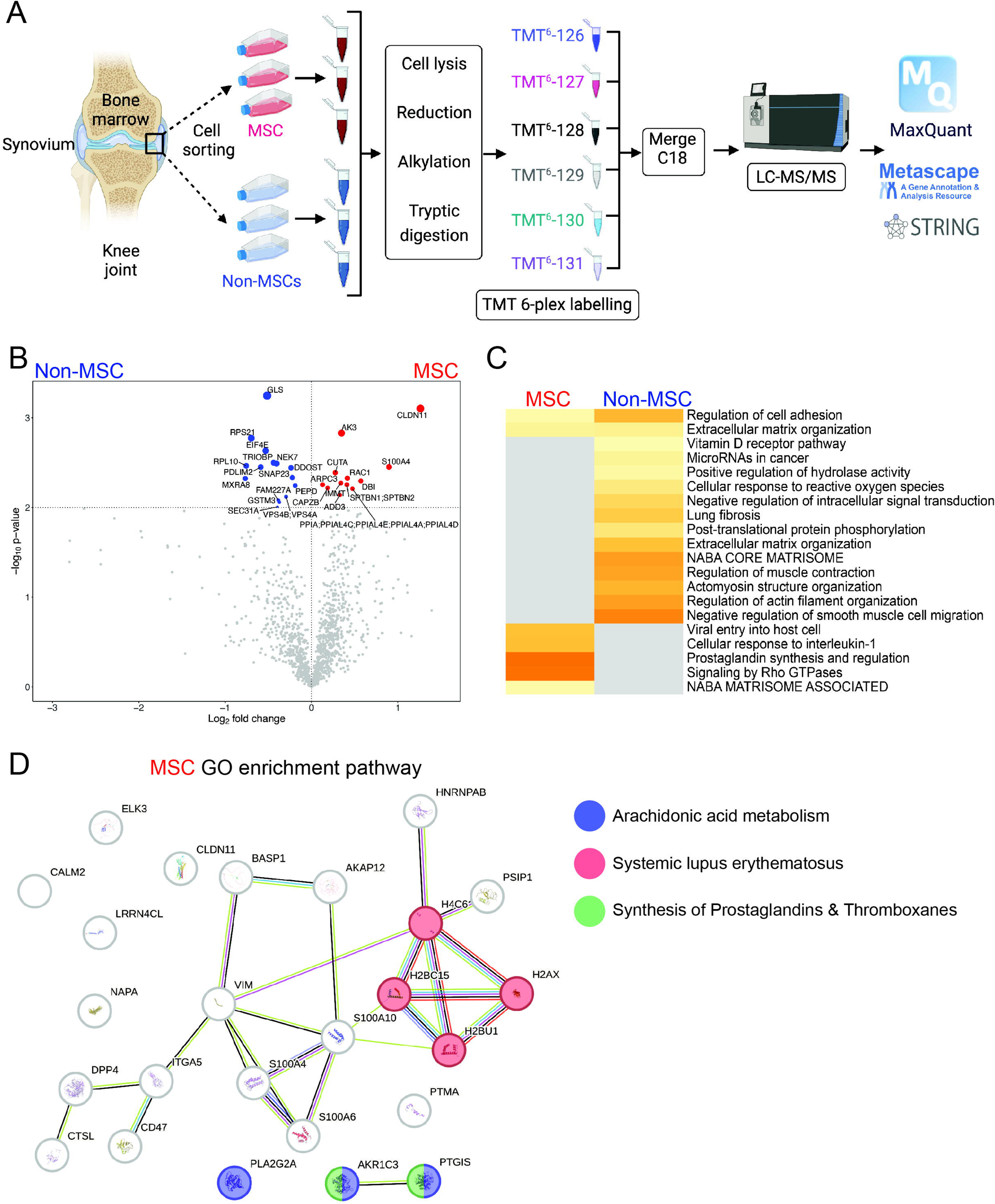
Proteomics analysis of MSC vs. non-MSC populations. Schematic representation of the quantitative shotgun proteomics workflow (A). Volcano plot demonstrating the distribution of statistically changing proteins identified in MSCs (red) and Non-MSCs (blue) (C). Metascape analysis and heatmap showing pathways regulated by differentially expressed proteins in MSCs vs. non-MSCs (D). STRING-db analysis of protein-protein interaction networks with elevated proteins in MSCs (D).

### CD47 as a marker of human synovial cells with in vivo chondrogenic potential

Synovial biopsies were recovered from an additional n=3 normal and n=3 OA knee joints and the CD90^+^CD73^+^CD44^+^ population was identified (**Figure 6A**). This population of cells was further interrogated for the expression of CD47, and it was found that all CD73^+^CD44^+^ double positive cells expressed CD47, but there was also a CD47^Hi^ population (**Figure 6B,C**) in both normal and OA synovium. The CD47^Hi^ and CD47^Lo^ sub-populations were both isolated by FACS and we further examined the immunophenotype of the cells. Specifically, we assayed if these CD47^Hi^ and CD47^Lo^ sub-populations expressed ITGA5 and/or DPP4/CD26 as these markers have previously been associated with stromal progenitors and shown to play a role in the progression of arthritis (Zheng et al., 2025). The CD47^Hi^ sub-population from normal and OA synovium expressed both ITGA5 and DPP4/CD26 whereas the CD47^Lo^ sub-population was largely negative for both markers (**Figure S7**). To further understand if there were any functional differences between sub-populations, we differentiated the cells into chondrocytes using standard pellet culture. Both CD47^Hi^ and CD47^Lo^ sub-populations produced pellets that stained positive for Alcian blue (**Figure 6 B,D**), however, the CD47^Hi^ cells appeared to have more instance staining. Therefore, we quantified the GAG content and found that CD47^Hi^ cells produced more GAGs that CD47^Lo^ cells regardless of if they were isolated from normal of OA synovium (**Figure 6 E**). Collagen 2 staining of the pellets showed a similar trend in where more pronounced Col2 staining was observed in pellets derived from CD47^Hi^ cells (**Figure 6 F,G**).

**Figure 6.**
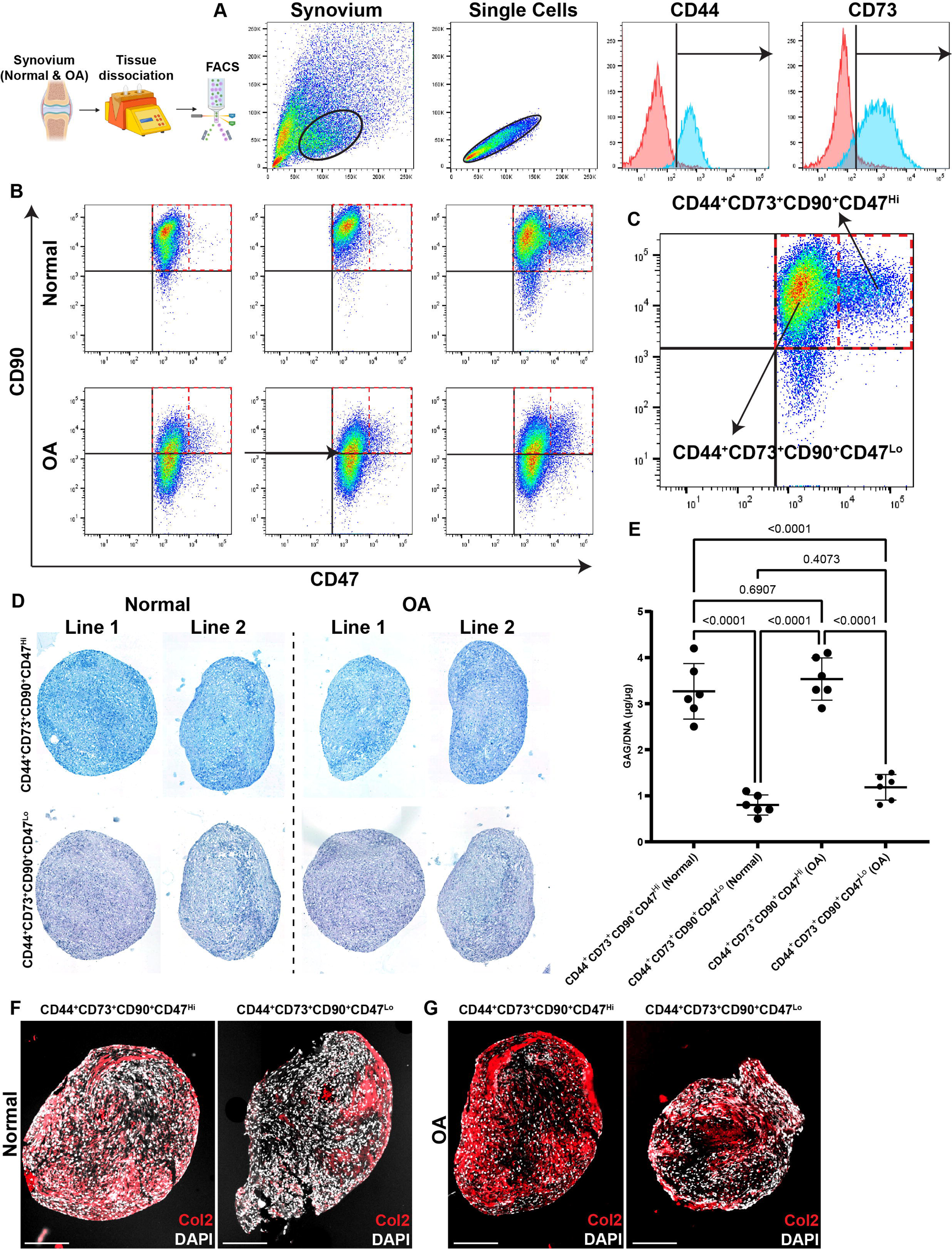
Analysis of synovial cells expressing CD47. The gating strategy to identify and sort CD47^Hi^ vs. CD47^Lo^ cell populations (A-C). CD47^Hi^ vs. CD47^Lo^ cells were identified and isolated from normal and OA synovial tissue (B). Chondrogenic differentiation of CD47^Hi^ vs. CD47^Lo^ cells using pellet culture and stained with Alican Blue (D). GAG quantification of the pellets (E). The pellets were also stained with Collagen 2 (Col2) as a marker of mature cartilage ECM (E,G). Scale bars equal 100µm.

These CD47^Hi^ and CD47^Lo^ populations were then injected into rats that had underwent DMM injury one week prior (**Figure 7 A**). Rats that received CD47^Hi^ cells presented with significantly lowers OARSI scores than rats receiving CD47^Lo^ cells (**Figure 7 B-D**), but there was no difference in Krenn scoring between rats which received CD47^Hi^ or CD47^Lo^ cells (**Figure 7 B**). When the joints tissues were isolated for histology, it was found that CD47^Hi^ cells contributed to the formation of new articular cartilage tissue within the injured joints (**Figure 7 H-J**), however, this was not observed with CD47^Lo^ cells (**Figure 7 E-G**). Specifically, human cells (tdTomato^+^) were observed within the cartilage of CD47^Hi^ injected rats, and these cells also expressed the chondrogenic markers Sox9 and Prg4 (**Figure 7 H-J**). In rats that received CD47^Lo^ cells, few to no tdTomato cells were observed within the injured cartilage (**Figure 7 E-G**).

**Figure 7.**
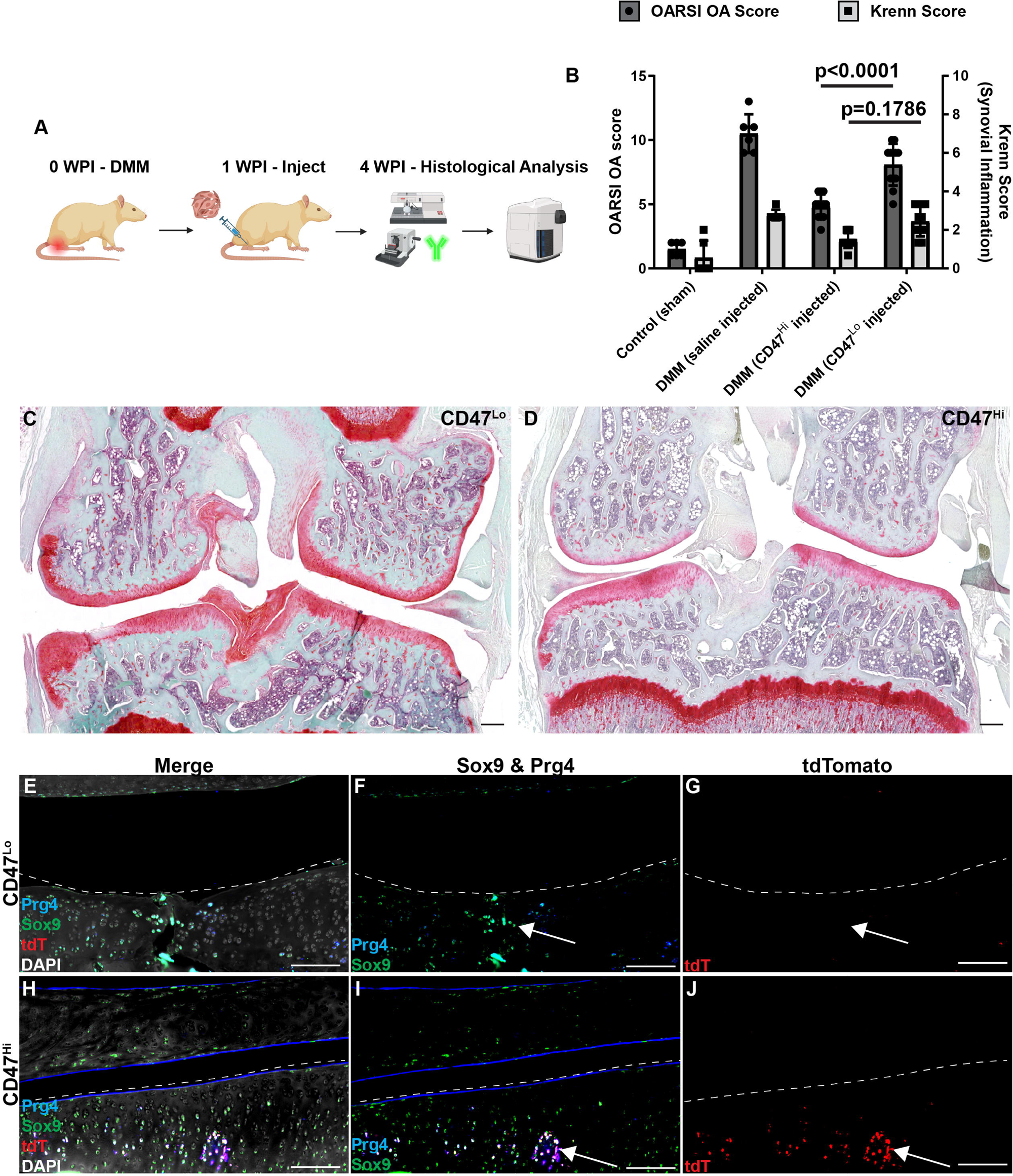
CD47^Hi^ vs. CD47^Lo^ cell contribution to cartilage repair in a rat DMM model. The CD47^Hi^ vs. CD47^Lo^ cells were injected into rats that underwent DMM injury (A) and it was observed that rat which received the CD47^Hi^ cells presented with significantly lower OARSI scores than rats receiving CD47^Lo^ cells (B) with representative images presented of joints receiving CD47^Hi^ (C) and CD47^Lo^ (D) cells. Immunofluorescence analysis demonstrated that the transplanted human CD47^Hi^ cells (TdTomato) integrated into the rat articular cartilage and expressed the chondrocyte markers Prg4 and Sox9 (H-J). This was not observed in the cartilage of rats that received the CD47^Lo^ cells (E-G). Scale bars equal 50µm.

## Discussion

The synovium plays a vital role in the overall health of the joint (Berenbaum, 2013; Kapoor et al., 2011). In a normal, healthy joint, synoviocytes assist in nourishing chondrocytes, removing waste and debris, and facilitating gas exchange. However, in OA, the breakdown of the cartilaginous matrix and subsequent phagocytosis of debris by synoviocytes promotes synovitis. This inflammation triggers the release of soluble proinflammatory mediators from the synovium, enhancing cartilage breakdown and perpetuating a vicious cycle. Although OA disrupts the primary function of articular cartilage (to provide cushioning and decrease friction within the joint), it impacts all tissues within the joint environment. The loss of the protective cartilage layer leads to intense pain in affected individuals. Moreover, damaged articular cartilage has limited ability to repair itself due to its avascular and aneural nature (Masson and Krawetz, 2020). Current non-surgical treatment strategies for OA focus on alleviating symptoms such as pain and inflammation. However, there are currently no proven treatments to prevent cartilage loss and promote tissue regrowth. While MSCs from the synovial membrane and synovial fluid have shown tri-lineage differentiation potential in various studies, endogenous articular cartilage repair remains elusive (De Bari, 2015; I et al., 2021; Krawetz et al., 2012; Masson et al., 2015; Sekiya et al., 2012). Moreover, distinct sub-populations of synovial MSCs exhibit widely different characteristics, including cell surface protein expression and differentiation potential(Affan et al., 2019). This suggests that not all tissue resident stem cells are created equal and underscores the need for further research into understanding these sub-populations.

The aim of this project was to characterize different progenitor populations in the synovium of normal and OA knee joints based on their self-renewal capacity, differentiation ability, and cell surface marker profile. However, relying solely on cell surface markers for MSC selection can be unreliable, as not all subpopulations demonstrate a high correlation between marker expression and differentiation potential. Moreover, changes in marker expression under different laboratory conditions can confound MSC characterization. Furthermore, it has been shown that some cell surface markers (such as CD105) may be an artifact of *in vitro* culturing as this molecule doesn’t appear to be expressed by MSCs *in vivo* and is in part responsible for the ability of MSCs to adhere to plastic (Anderson et al., 2013; Cleary et al., 2016). Therefore, our study utilized single-cell sorting to isolate cells from human knee samples, minimizing exposure to the culture microenvironment before analysis. We found that regardless of *in situ* marker expression, nearly all clonal cell populations displayed a uniform marker profile consistent with a presumptive MSC phenotype. This strongly suggests that the *in vitro* micro-environment upregulates and/or stabilizes the expression of the receptors. This notion is supported by our observation (similar to previous published studies)(Affan et al., 2019), that the expression of CD105 was acquired only after culture, indicating its potential role as a bystander marker in MSC characterization. Additionally, the expression of CD271 varied among MSC clones, with none of the clones demonstrating multi-potential differentiation capacity expressing CD271. This suggests that CD271 may not be a marker of synovial MSCs(Barilani et al., 2018). Also interesting, was that the distribution of OA MSC subpopulations differed from those in healthy joints, indicating distinct alterations in the synovial niche during OA onset/progression. In future studies, it would be interesting to determine what exact aspects of *in vitro* culture cause the cells to take on a uniform immunophenotype. Specifically, testing out different media and/or matrix compositions could shed light on this observation.

In the current study, we have identified a subpopulation of synovial cells that express high levels of CD47. Our findings support the hypothesis that cells expressing high levels of CD47 have robust cartilage regeneration potential. CD47, also known as integrin-associated protein, is a transmembrane protein that plays a pivotal role in immune regulation. It acts as a “don’t eat me” signal, inhibiting phagocytosis by macrophages and other immune cells(Hao et al., 2023; Maute et al., 2022; Oldenborg, 2013). It has been shown that overexpression of CD47 allows cells (such as in various cancers) to evade immune surveillance. In stem cell populations, particularly adult stem cells, CD47 plays a crucial role in maintaining self-renewal and tissue regeneration capacities (Jaiswal et al., 2009; Kim et al., 2021). It regulates stem cell survival and proliferation by interacting with its receptor, SIRPα, on neighboring cells or stromal components. This interaction influences stem cell behavior, homing, and differentiation, shaping tissue homeostasis and repair processes. Its upregulation on dying cells can lead to the inhibition of phagocytosis of these necrotic/apoptotic cells (Kojima et al., 2016), leading to the accumulation of cellular debris within the environment, thus perpetuating inflammation. Overall, there is still not much known about how CD47 regulates stem cell/progenitor function, and it would be of interest to experimentally test if the overexpression of CD47 in MSCs can increase their endogenous cartilage regeneration capacity.

While the clonal derivation and long-term expansion of cells made this experimental design possible, this approach is not without limitations. Firstly, during clonal derivation, cells may acquire genetic mutations or epigenetic alterations due to the stresses of isolation and culture conditions, leading to phenotypic changes and potential change/loss of functionality. Moreover, the process of long-term expansion *in vitro* can induce replicative senescence, telomere shortening, and genomic instability, compromising the quality and integrity of the cell population over time. This could account in part for the majority of clones we derived losing the ability to proliferate before we could achieve a sufficient number of cells for the downstream analyses. Furthermore, cells cultured *in vitro* often face challenges in recapitulating the complex microenvironment and dynamic interactions present with their *in vivo* niche(s), potentially altering their behavior and functional properties. As such, the reliance on exogenous growth factors, culture media, and artificial substrates to support cell growth and expansion introduces variability and may not fully mimic the physiological cues present in the native tissue environment. Such discrepancies can impact the reliability and reproducibility of experimental outcomes and may hinder the successful clinical translation of cell-based therapies. Yet, even with these challenges, we have presented compelling evidence demonstrating that a sub-population of cells exist within the human synovial membrane (under normal and OA conditions) that can contribute to articular cartilage regeneration.

In conclusion, while MSCs hold promise for OA treatment, the heterogeneity within MSC populations presents challenges for their effective utilization. The development of effective and reproducible protocols for MSC identification/characterization are necessary to ensure the selection of MSCs with the greatest potential for joint repair and regeneration. With this knowledge, we have the best chance of developing novel cell therapies that will benefit the growing number of patients suffering from OA.

## Acknowledgements

The authors would like to thank Dr. Yiping Liu of the Cumming School of Medicine Flow Cytometry Facility for assistance with FACS and the Cumming School of Medicine ARC staff for assistance with animal husbandry.

## Author Contributions

Conception and design: NAJ, RJK. Analysis and interpretation of the data: NAJ, AA, CL, LGA, AD, PS, RJK. Drafting of the article: NAJ, RJK. Critical revision of the article for important intellectual content: NAJ, AA, CL, ND, LGA, DY, AOM, AD, PS, PM, JNP, RJK. Provision of study materials: PR, JNP, RJK. Obtaining of funding: RJK. Collection and assembly of data: NAJ, AD, RJK. Final approval of the article for submission: NAJ, AA, CL, ND, LGA, DY, AOM, AD, PS, PM, JNP, RJK.

## Role of the funding source

Natural Sciences and Engineering Research Council (NSERC) of Canada RGPIN-2014-04586 (RJK). Canada Foundation for Innovation (RJK). Calgary Foundation, Grace Glaum Professorship (RJK). The funders had no role in study design, data collection and analysis, decision to publish, or preparation of the manuscript.

## Competing interest statement

The authors declare no competing interests.

## Data availability statement

Data available within the article or its supplementary materials.

## Notes

### Competing Interest Statement

The authors have declared no competing interest.

### Summary of Updates

Revisions have been made to address the comments and concerns of the reviewers.

## References

1. Affan A, Al-Jezani N, Railton P, Powell JN, Krawetz RJ. 2019. Multiple mesenchymal progenitor cell subtypes with distinct functional potential are present within the intimal layer of the hip synovium. BMC Musculoskelet Disord 20:125. doi:10.1186/s12891-019-2495-2

2. Anderson P, Carrillo-Gálvez AB, García-Pérez A, Cobo M, Martín F. 2013. CD105 (endoglin)-negative murine mesenchymal stromal cells define a new multipotent subpopulation with distinct differentiation and immunomodulatory capacities. PLoS One 8:e76979. doi:10.1371/journal.pone.0076979

3. Arden NK, Perry TA, Bannuru RR, Bruyère O, Cooper C, Haugen IK, Hochberg MC, McAlindon TE, Mobasheri A, Reginster J-Y. 2020. Non-surgical management of knee osteoarthritis: comparison of ESCEO and OARSI 2019 guidelines. Nat Rev Rheumatol. doi:10.1038/s41584-020-00523-9

4. Barilani M, Banfi F, Sironi S, Ragni E, Guillaumin S, Polveraccio F, Rosso L, Moro M, Astori G, Pozzobon M, Lazzari L. 2018. Low-affinity Nerve Growth Factor Receptor (CD271) Heterogeneous Expression in Adult and Fetal Mesenchymal Stromal Cells. Sci Rep 8:9321. doi:10.1038/s41598-018-27587-8

5. Berenbaum F. 2013. Osteoarthritis as an inflammatory disease (osteoarthritis is not osteoarthrosis!). Osteoarthr Cartil 21:16–21. doi:10.1016/j.joca.2012.11.012

6. Cleary MA, Narcisi R, Focke K, van der Linden R, Brama PAJ, van Osch GJVM. 2016. Expression of CD105 on expanded mesenchymal stem cells does not predict their chondrogenic potential. Osteoarthr Cartil 24:868–872. doi:10.1016/j.joca.2015.11.018

7. Collins FL, Roelofs AJ, Symons RA, Kania K, Campbell E, Collie-Duguid ESR, Riemen AHK, Clark SM, De Bari C. 2023. Taxonomy of fibroblasts and progenitors in the synovial joint at single-cell resolution. Ann Rheum Dis 82:428–437. doi:10.1136/ard-2021-221682

8. Das N, de Almeida LGN, Derakhshani A, Young D, Mehdinejadiani K, Salo P, Rezansoff A, Jay GD, Sommerhoff CP, Schmidt TA, Krawetz R, Dufour A. 2023. Tryptase β regulation of joint lubrication and inflammation via proteoglycan-4 in osteoarthritis. Nat Commun 14:1910. doi:10.1038/S41467-023-37598-3

9. Das N, Schmidt TA, Krawetz RJ, Dufour A. 2019. Proteoglycan 4: From Mere Lubricant to Regulator of Tissue Homeostasis and Inflammation. BioEssays 41:1800166. doi:10.1002/bies.201800166

10. De Bari C. 2015. Are mesenchymal stem cells in rheumatoid arthritis the good or bad guys? Arthritis Res Ther 17:113. doi:10.1186/s13075-015-0634-1

11. De Bari C, Dell’Accio F, Tylzanowski P, Luyten FP. 2001. Multipotent mesenchymal stem cells from adult human synovial membrane. Arthritis Rheum 44:1928–1942. doi:10.1002/1529-0131(200108)44:8<1928::AID-ART331>3.0.CO;2-P

12. de Witte SFH, Luk F, Sierra Parraga JM, Gargesha M, Merino A, Korevaar SS, Shankar AS, O’Flynn L, Elliman SJ, Roy D, Betjes MGH, Newsome PN, Baan CC, Hoogduijn MJ. 2018. Immunomodulation By Therapeutic Mesenchymal Stromal Cells (MSC) Is Triggered Through Phagocytosis of MSC By Monocytic Cells. Stem Cells 36:602–615. doi:10.1002/stem.2779

13. Deshmane SL, Kremlev S, Amini S, Sawaya BE. 2009. Monocyte chemoattractant protein-1 (MCP-1): an overview. J Interferon Cytokine Res 29:313–26. doi:10.1089/jir.2008.0027

14. DiMarino AM, Caplan AI, Bonfield TL. 2013. Mesenchymal stem cells in tissue repair. Front Immunol. doi:10.3389/fimmu.2013.00201

15. Dominici M, Le Blanc K, Mueller I, Slaper-Cortenbach I, Marini F. C, Krause DSS, Deans RJJ, Keating A, Prockop DJJ, Horwitz EMM. 2006. Minimal criteria for defining multipotent mesenchymal stromal cells. The International Society for Cellular Therapy position statement. Cytotherapy 8:315–317. doi:10.1080/14653240600855905

16. Fellows CR, Matta C, Zakany R, Khan IM, Mobasheri A. 2016. Adipose, Bone Marrow and Synovial Joint-Derived Mesenchymal Stem Cells for Cartilage Repair. Front Genet 7:213. doi:10.3389/fgene.2016.00213

17. Fernandes TL, Kimura HA, Pinheiro CCG, Shimomura K, Nakamura N, Ferreira JR, Gomoll AH, Hernandez AJ, Bueno DF. 2018. Human synovial mesenchymal stem cells good manufacturing practices for articular cartilage regeneration. Tissue Eng – Part C Methods 24:709–716. doi:10.1089/ten.tec.2018.0219

18. Futami I, Ishijima M, Kaneko H, Tsuji K, Ichikawa-Tomikawa N, Sadatsuki R, Muneta T, Arikawa-Hirasawa E, Sekiya I, Kaneko K. 2012. Isolation and Characterization of Multipotential Mesenchymal Cells from the Mouse Synovium. PLoS One 7. doi:10.1371/journal.pone.0045517

19. Grandi FC, Bhutani N. 2020. Epigenetic Therapies for Osteoarthritis. Trends Pharmacol Sci 41:557–569. doi:10.1016/j.tips.2020.05.008

20. Hafsi K, McKay J, Li J, Lana JF, Macedo A, Santos GS, Murrell WD. 2019. Nutritional, metabolic and genetic considerations to optimise regenerative medicine outcome for knee osteoarthritis. J Clin Orthop Trauma 10:2–8. doi:10.1016/j.jcot.2018.10.004

21. Hao Y, Zhou X, Li Y, Li B, Cheng L. 2023. The CD47-SIRPα axis is a promising target for cancer immunotherapies. Int Immunopharmacol 120. doi:10.1016/j.intimp.2023.110255

22. Harris Q, Seto J, O’Brien K, Lee PS, Kondo C, Heard BJ, Hart DA, Krawetz RJ. 2013. Monocyte chemotactic protein-1 inhibits chondrogenesis of synovial mesenchymal progenitor cells: an in vitro study. Stem Cells 31:2253–65. doi:10.1002/stem.1477

23. I S, H K, N O. 2021. Characteristics of MSCs in Synovial Fluid and Mode of Action of Intra-Articular Injections of Synovial MSCs in Knee Osteoarthritis. Int J Mol Sci 22:1–13. doi:10.3390/IJMS22062838

24. Iqbal SM, Leonard C, Regmi SC, De Rantere D, Tailor P, Ren G, Ishida H, Hsu C, Abubacker S, Pang DS, Salo PT, Vogel HJ, Hart DA, Waterhouse CC, Jay GD, Schmidt TA, Krawetz RJ. 2016. Lubricin/Proteoglycan 4 binds to and regulates the activity of Toll-Like Receptors In Vitro. Sci Rep 6:18910. doi:10.1038/srep18910

25. Jablonski CL, Leonard C, Salo P, Krawetz RJ. 2019. CCL2 But Not CCR2 Is Required for Spontaneous Articular Cartilage Regeneration Post-Injury. J Orthop Res 37:2561–2574. doi:10.1002/jor.24444

26. Jaiswal S, Jamieson CHM, Pang WW, Park CY, Chao MP, Majeti R, Traver D, van Rooijen N, Weissman IL. 2009. CD47 is upregulated on circulating hematopoietic stem cells and leukemia cells to avoid phagocytosis. Cell 138:271–285. doi:10.1016/J.CELL.2009.05.046

27. Jia Z, Liu Q, Liang Y, Li X, Xu X, Ouyang K, Xiong J, Wang D, Duan L. 2018. Repair of articular cartilage defects with intra-articular injection of autologous rabbit synovial fluid-derived mesenchymal stem cells. J Transl Med 16:1–12. doi:10.1186/s12967-018-1485-8

28. Kapoor M, Martel-Pelletier J, Lajeunesse D, Pelletier J-P, Fahmi H. 2011. Role of proinflammatory cytokines in the pathophysiology of osteoarthritis. Nat Rev Rheumatol 7:33–42. doi:10.1038/nrrheum.2010.196

29. Kim GH, Bae YK, Kwon JH, Kim M, Choi SJ, Oh W, Um S, Jin HJ. 2021. Positively Correlated CD47 Activation and Autophagy in Umbilical Cord Blood-Derived Mesenchymal Stem Cells during Senescence. Stem Cells Int 2021. doi:10.1155/2021/5582792

30. Kim JH, Jo CH, Kim HR, Hwang Y Il. 2018. Comparison of immunological characteristics of mesenchymal stem cells from the periodontal ligament, umbilical cord, and adipose tissue. Stem Cells Int 2018. doi:10.1155/2018/8429042

31. Knights AJ, Farrell EC, Ellis OM, Lammlin L, Junginger LM, Rzeczycki PM, Bergman RF, Pervez R, Cruz M, Knight E, Farmer D, Samani AA, Wu CL, Hankenson KD, Maerz T. 2022. Synovial fibroblasts assume distinct functional identities and secrete R-spondin 2 in osteoarthritis. Ann Rheum Dis 82:272–282. doi:10.1136/ard-2022-222773

32. Koh YG, Kwon OR, Kim YS, Choi YJ, Tak DH. 2016. Adipose-Derived Mesenchymal Stem Cells With Microfracture Versus Microfracture Alone: 2-Year Follow-up of a Prospective Randomized Trial. Arthroscopy 32:97–109. doi:10.1016/J.ARTHRO.2015.09.010

33. Kojima Y, Volkmer JP, McKenna K, Civelek M, Lusis AJ, Miller CL, Direnzo D, Nanda V, Ye J, Connolly AJ, Schadt EE, Quertermous T, Betancur P, Maegdefessel L, Matic LP, Hedin U, Weissman IL, Leeper NJ. 2016. CD47-blocking antibodies restore phagocytosis and prevent atherosclerosis. Nature 536:86–90. doi:10.1038/NATURE18935

34. Krawetz RJ, Affan A, Leonard C, Veeramreddy DN, Fichadiya A, Martin L, Schmeling H. 2022. Synovial fluid mesenchymal progenitor cells from patients with juvenile idiopathic arthritis demonstrate limited self-renewal and chondrogenesis. Sci Rep 12. doi:10.1038/S41598-022-20880-7

35. Krawetz RJ, Larijani L, Corpuz JM, Ninkovic N, Das N, Olsen A, Mohtadi N, Rezansoff A, Dufour A. 2023. Mesenchymal progenitor cells from non-inflamed versus inflamed synovium post-ACL injury present with distinct phenotypes and cartilage regeneration capacity. Stem Cell Res Ther 14:168. doi:10.1186/s13287-023-03396-3

36. Krawetz RJ, Wu YE, Martin L, Rattner JB, Matyas JR, Hart DA. 2012. Synovial Fluid Progenitors Expressing CD90+ from Normal but Not Osteoarthritic Joints Undergo Chondrogenic Differentiation without Micro-Mass Culture. PLoS One 7. doi:10.1371/journal.pone.0043616

37. Lee WJ, Hah YS, Ock SA, Lee JH, Jeon RH, Park JS, Lee S Il, Rho NY, Rho GJ, Lee SL. 2015. Cell source-dependent in vivo immunosuppressive properties of mesenchymal stem cells derived from the bone marrow and synovial fluid of minipigs. Exp Cell Res 333:273–288. doi:10.1016/j.yexcr.2015.03.015

38. Li J, Gui T, Yao L, Guo H, Lin YL, Lu J, Duffy M, Zgonis M, Mauck R, Dyment N, Zhang Y, Scanzello C, Seale P, Qin L. 2024. Synovium and infrapatellar fat pad share common mesenchymal progenitors and undergo coordinated changes in osteoarthritis. J Bone Miner Res 39:161–176. doi:10.1093/JBMR/ZJAD009,

39. Mak J, Jablonski CL, Leonard CA, Dunn JF, Raharjo E, Matyas JR, Biernaskie J, Krawetz RJ. 2016. Intra-articular injection of synovial mesenchymal stem cells improves cartilage repair in a mouse injury model. Sci Rep 6. doi:10.1038/srep23076

40. Masson AO, Hess R, O’Brien K, Bertram KL, Tailor P, Irvine E, Ren G, Krawetz RJ. 2015. Increased levels of p21(CIP1/WAF1) correlate with decreased chondrogenic differentiation potential in synovial membrane progenitor cells. Mech Ageing Dev 149. doi:10.1016/j.mad.2015.05.005

41. Masson AO, Krawetz RJ. 2020. Understanding cartilage protection in OA and injury: a spectrum of possibilities. BMC Musculoskelet Disord 21. doi:10.1186/S12891-020-03363-6

42. Maute R, Xu J, Weissman IL. 2022. CD47-SIRPα-targeted therapeutics: status and prospects. Immuno-oncology Technol 13. doi:10.1016/J.IOTECH.2022.100070

43. Mautner K, Gottschalk M, Boden SD, Akard A, Bae WC, Black L, Boggess B, Chatterjee P, Chung CB, Easley KA, Gibson G, Hackel J, Jensen K, Kippner L, Kurtenbach C, Kurtzberg J, Mason RA, Noonan B, Roy K, Valentine V, Yeago C, Drissi H. 2023. Cell-based versus corticosteroid injections for knee pain in osteoarthritis: a randomized phase 3 trial. Nat Med 2023 2912 29:3120–3126. doi:10.1038/s41591-023-02632-w

44. Miller RE, Tran PB, Das R, Ghoreishi-Haack N, Ren D, Miller RJ, Malfait A-M. 2012. CCR2 chemokine receptor signaling mediates pain in experimental osteoarthritis. Proc Natl Acad Sci U S A 109:20602–7. doi:10.1073/pnas.1209294110

45. Mora JC, Przkora R, Cruz-Almeida Y. 2018. Knee osteoarthritis: pathophysiology and current treatment modalities. J Pain Res Volume 11:2189–2196. doi:10.2147/JPR.S154002

46. Ni Z, Zhou S, Li S, Kuang L, Chen H, Luo X, Ouyang J, He M, Du X, Chen L. 2020. Exosomes: roles and therapeutic potential in osteoarthritis. Bone Res 8:25. doi:10.1038/s41413-020-0100-9

47. Nishimura K, Solchaga LA, Caplan AI, Yoo JU, Goldberg VM, Johnstone B. 1999. Chondroprogenitor cells of synovial tissue. Arthritis Rheum 42:2631–2637. doi:10.1002/1529-0131(199912)42:12<2631::AID-ANR18>3.0.CO;2-H

48. Oldenborg P-A. 2013. CD47: A Cell Surface Glycoprotein Which Regulates Multiple Functions of Hematopoietic Cells in Health and Disease. ISRN Hematol 2013:1–19. doi:10.1155/2013/614619

49. Palazzo C, Ravaud JF, Papelard A, Ravaud P, Poiraudeau S. 2014. The burden of musculoskeletal conditions. PLoS One 9. doi:10.1371/journal.pone.0090633

50. Peters H, Potla P, Rockel JS, Tockovska T, Pastrello C, Jurisica I, Delos Santos K, Vohra S, Fine N, Lively S, Perry K, Looby N, Li SH, Chandran V, Hueniken K, Kaur P, Perruccio A V., Mahomed NN, Rampersaud R, Syed K, Gracey E, Krawetz R, Buechler MB, Gandhi R, Kapoor M. 2025. Cell and transcriptomic diversity of infrapatellar fat pad during knee osteoarthritis. Ann Rheum Dis 84:351–367. doi:10.1136/ard-2024-225928

51. Phinney DG. 2012. Functional heterogeneity of mesenchymal stem cells: implications for cell therapy. J Cell Biochem 113:2806–12. doi:10.1002/jcb.24166

52. Raghu H, Lepus CM, Wang Q, Wong HH, Lingampalli N, Oliviero F, Punzi L, Giori NJ, Goodman SB, Chu CR, Sokolove JB, Robinson WH. 2017. CCL2/CCR2, but not CCL5/CCR5, mediates monocyte recruitment, inflammation and cartilage destruction in osteoarthritis. Ann Rheum Dis 76:914–922. doi:10.1136/annrheumdis-2016-210426

53. Russell KC, Phinney DG, Lacey MR, Barrilleaux BL, Meyertholen KE, O’Connor KC. 2010. In vitro high-capacity assay to quantify the clonal heterogeneity in trilineage potential of mesenchymal stem cells reveals a complex hierarchy of lineage commitment. Stem Cells 28:788–98. doi:10.1002/stem.312

54. Sakaguchi Y, Sekiya I, Yagishita K, Muneta T. 2005. Comparison of human stem cells derived from various mesenchymal tissues: Superiority of synovium as a cell source. Arthritis Rheum 52:2521–2529. doi:10.1002/art.21212

55. Salmon JH, Rat AC, Sellam J, Michel M, Eschard JP, Guillemin F, Jolly D, Fautrel B. 2016. Economic impact of lower-limb osteoarthritis worldwide: a systematic review of cost-of-illness studies. Osteoarthr Cartil 24:1500–1508. doi:10.1016/j.joca.2016.03.012

56. Satué M, Schüler C, Ginner N, Erben RG. 2019. Intra-articularly injected mesenchymal stem cells promote cartilage regeneration, but do not permanently engraft in distant organs. Sci Rep 9:1–10. doi:10.1038/s41598-019-46554-5

57. Sekiya I, Ojima M, Suzuki S, Yamaga M, Horie M, Koga H, Tsuji K, Miyaguchi K, Ogishima S, Tanaka H, Muneta T. 2012. Human mesenchymal stem cells in synovial fluid increase in the knee with degenerated cartilage and osteoarthritis. J Orthop Res 30:943–9. doi:10.1002/jor.22029

58. Soland MA, Bego M, Colletti E, Zanjani ED, St. Jeor S, Porada CD, Almeida-Porada G. 2013. Mesenchymal stem cells engineered to inhibit complement-mediated damage. PLoS One 8. doi:10.1371/JOURNAL.PONE.0060461

59. Tang S, Yao L, Ruan J, Kang J, Cao Y, Nie X, Lan W, Zhu Z, Han W, Liu Y, Tian J, Seale P, Qin L, Ding C. 2024. Single-cell atlas of human infrapatellar fat pad and synovium implicates APOE signaling in osteoarthritis pathology. Sci Transl Med 16:4590. doi:10.1126/SCITRANSLMED.ADF4590,

60. Zheng L, Gu M, Li X, Hu X, Chen C, Kang Y, Pan B, Chen W, Xian G, Wu X, Li C, Wang C, Li Z, Guan M, Zhou G, Mobasheri A, Song W, Peng S, Sheng P, Zhang Z. 2025. ITGA5+synovial fibroblasts orchestrate proinflammatory niche formation by remodelling the local immune microenvironment in rheumatoid arthritis. Ann Rheum Dis 84:232–252. doi:10.1136/ard-2024-225778

61. Zhou W, Lin J, Zhao K, Jin K, He Q, Hu Y, Feng G, Cai Y, Xia C, Liu H, Shen W, Hu X, Ouyang H. 2019. Single-Cell Profiles and Clinically Useful Properties of Human Mesenchymal Stem Cells of Adipose and Bone Marrow Origin. Am J Sports Med 47:1722– 1733. doi:10.1177/0363546519848678

