## Supplemental Figures and Tables for "Identification of a sub-population of synovial mesenchymal stem cells with enhanced treatment efficacy in a rat model of Osteoarthritis"

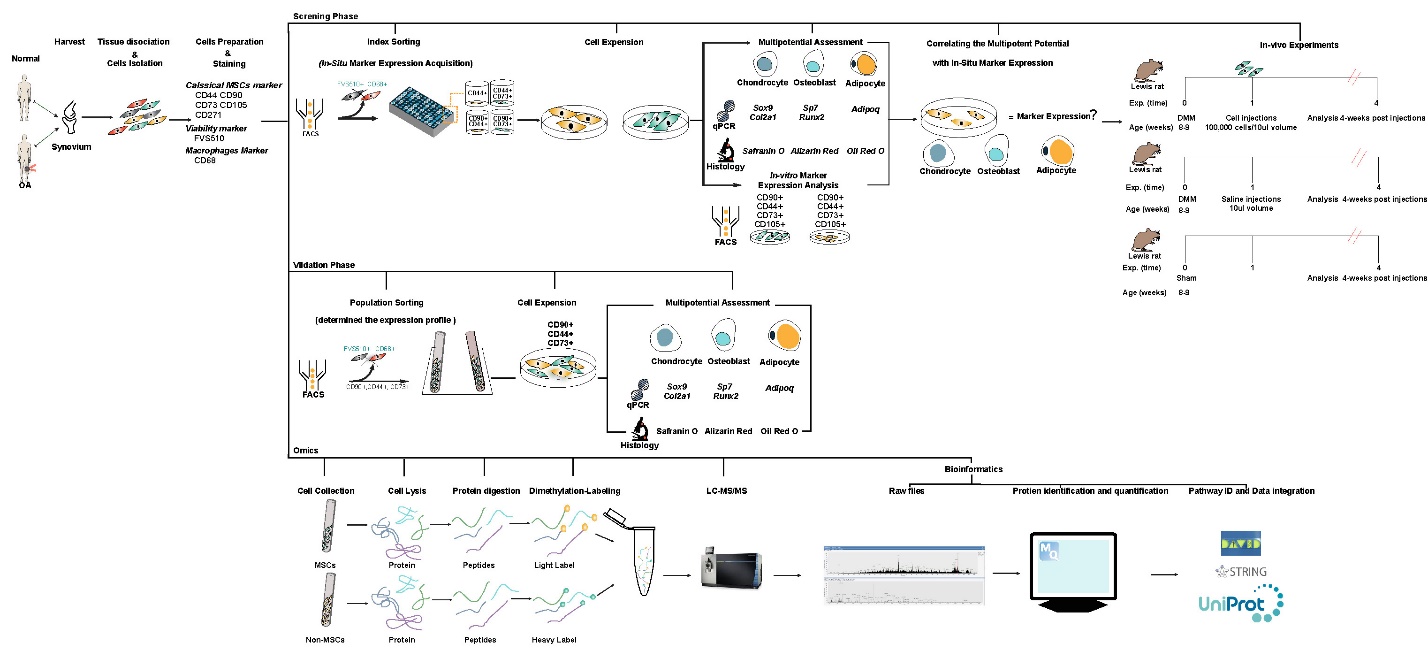

**Figure S1. Overview of the experimental design employed in the current study.**

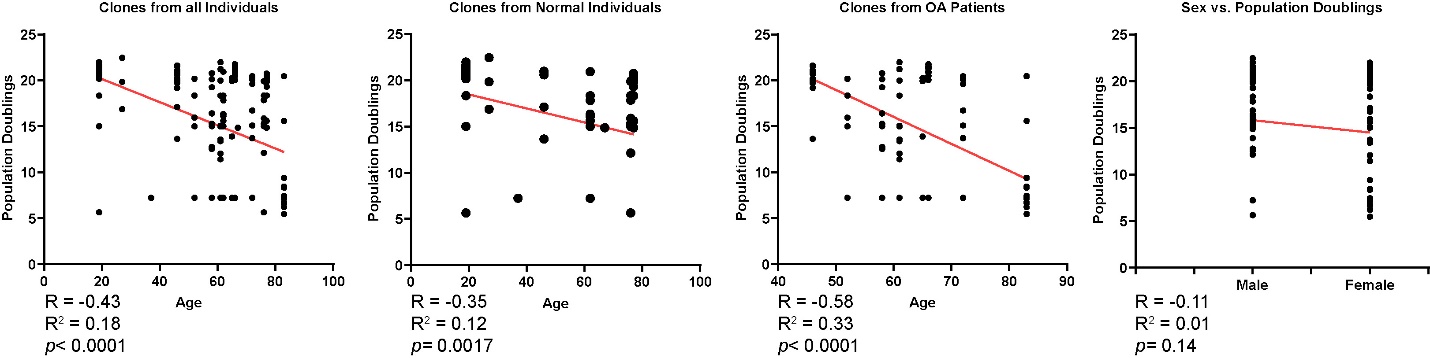

**Figure S2. Association between population doublings, sex and age.** A significant negative association was observed between age and population doublings in clones derived from normal and OA synovium. There was no association between sex and population doubling.

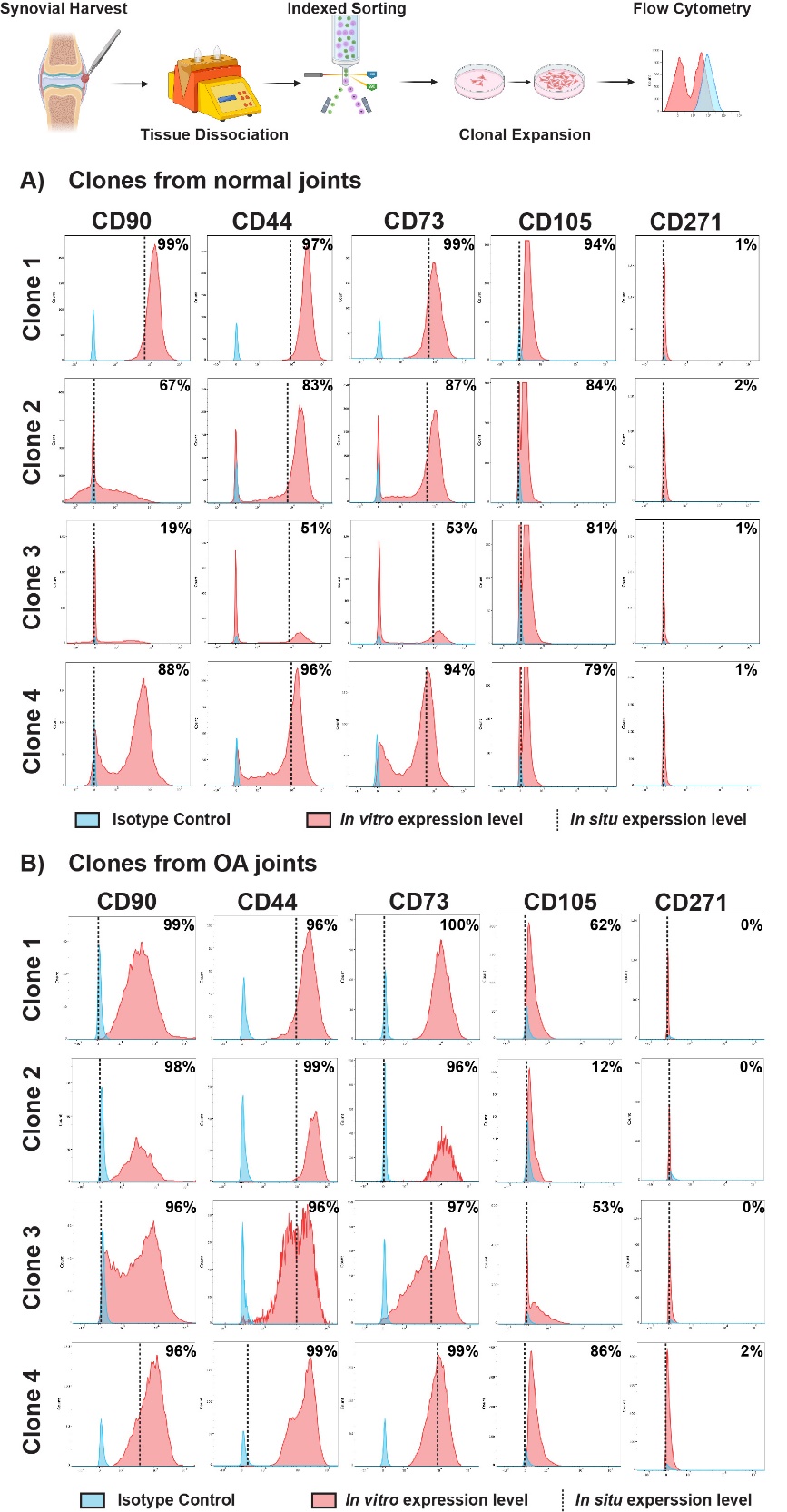

**Figure S3. *In situ* and *in vitro* cell surface protein expression on clones derived from normal individuals and patients with OA**. Representative histograms from 4 clones from normal (A) and OA (B) synovium are shown. The *in situ* expression of each CD marker is represented by the vertical dashed black bar. The *in vitro* expression of each marker in the clonal derived cell population is represented by the red histogram. The isotype/negative control for each CD marker is represented by the blue histogram.

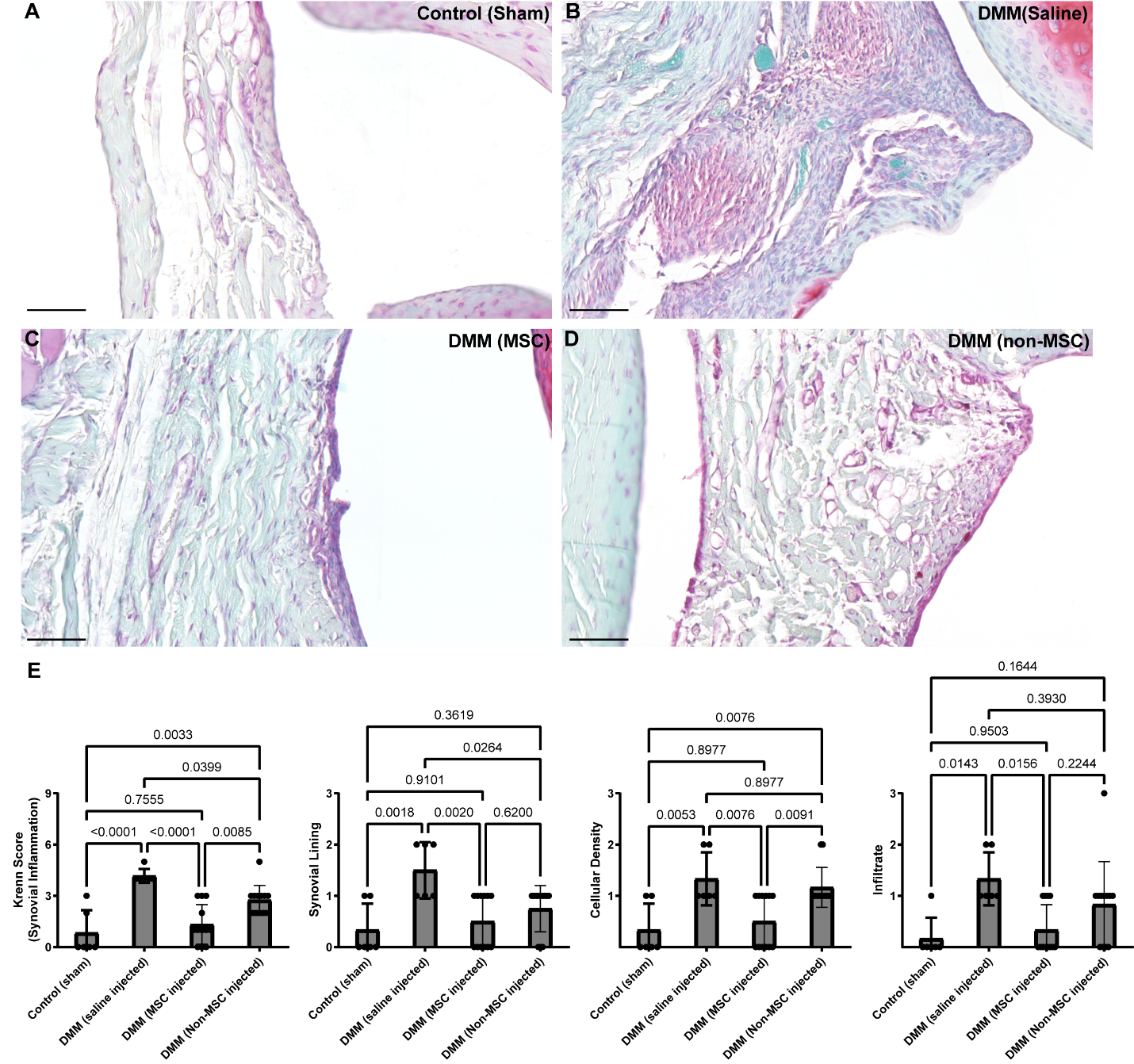

**Figure S4. Safranin O staining sections of rat synovium with or without cell delivery.** Representative images of synovium tissue from all four treatment groups (A-D). The summed Krenn score shown along with the individual categories including synovial lining, cellular density and infiltration (E). Scale bars equal 40µm.

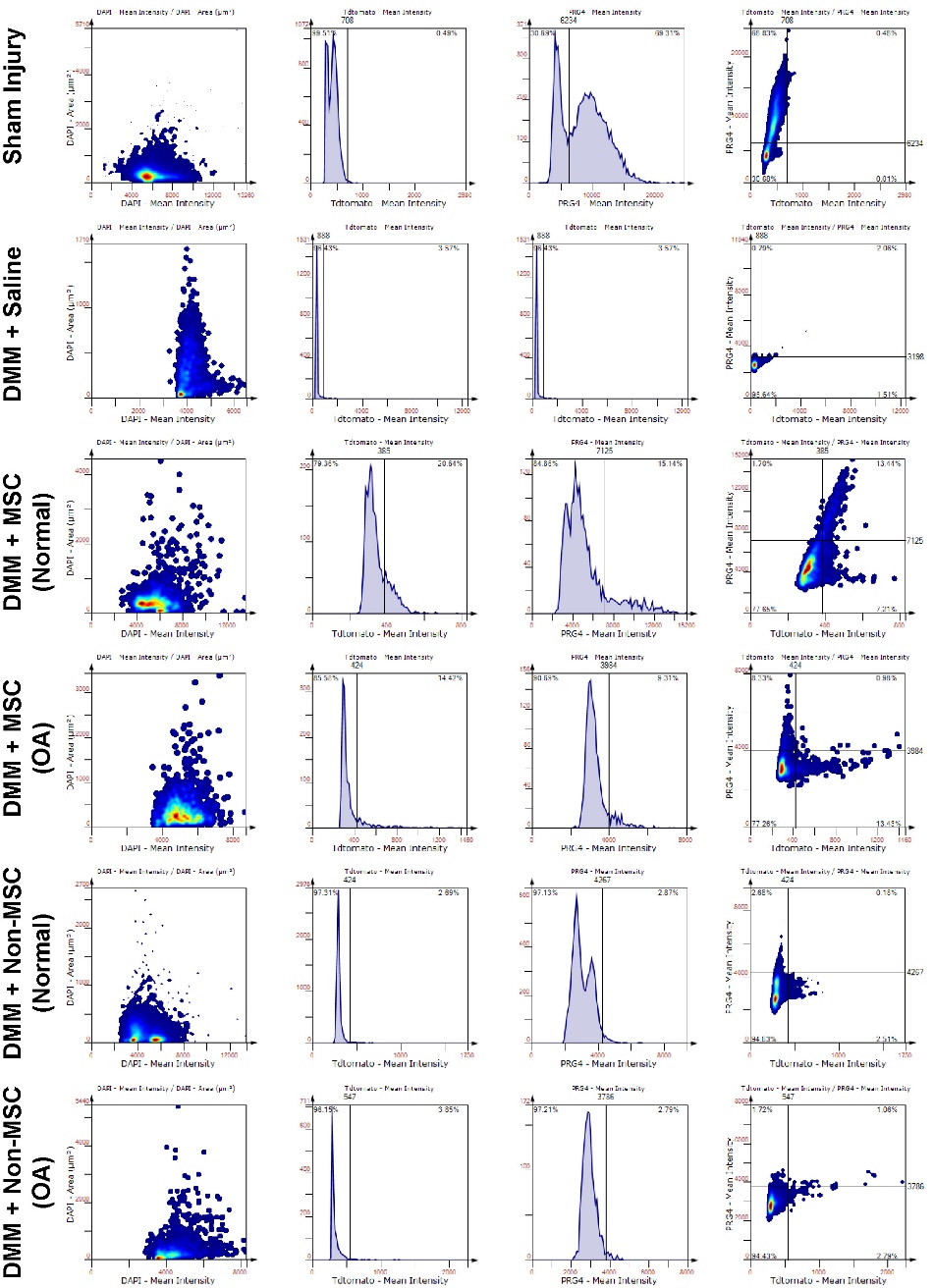

**Figure S5. Representative tissue cytometry gates**. Examples of gating strategy for each marker (DAPI, tdTomato, PRG4, tdTomato plus PRG4) in each experimental group.

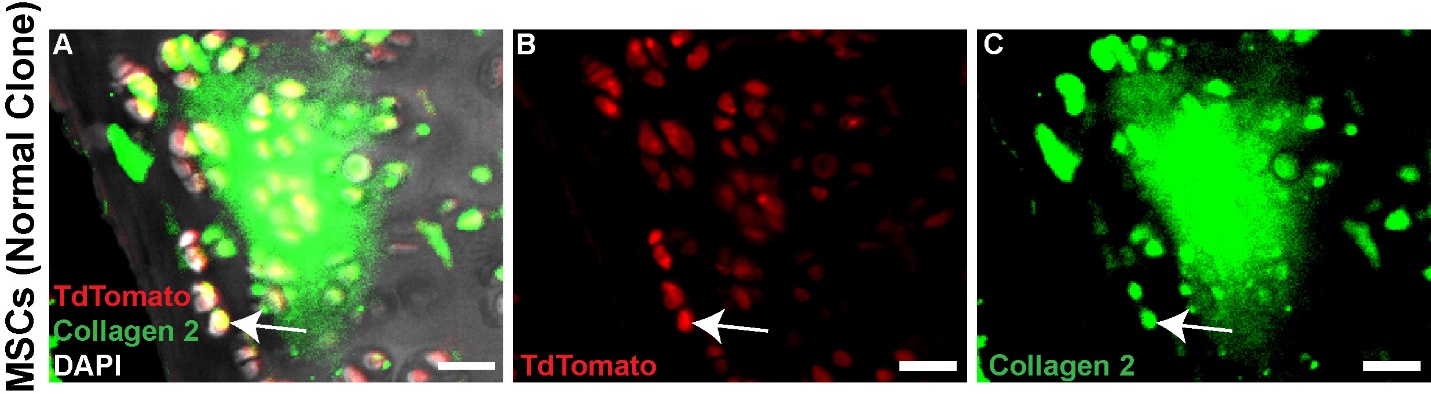

**Figure S6. MSCs within the cartilage produce collagen 2.** Representative images of a rat with DMM surgery (A) with MSC injection (TdTomato – B) and collagen 2 staining (C). Scale bars equal 20µm.

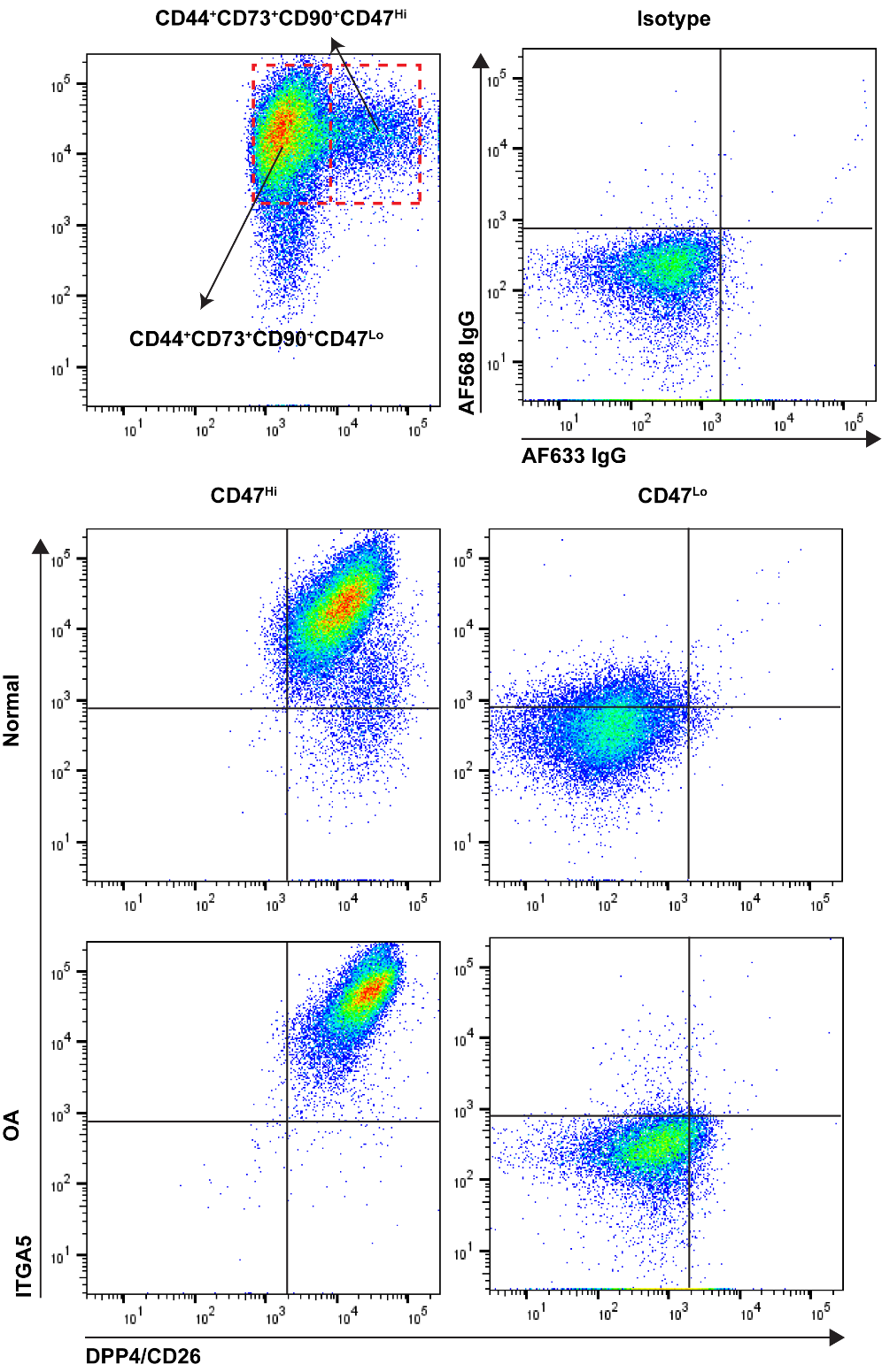

**Figure S7. Analysis of synovial cells expressing CD47**. CD47 cells from normal and OA synovium were assayed for the expression of ITAG5 and DPP4/CD26 by flow cytometry. The isotype control shows little to no reactivity. CD47^Hi^ cells co-expressed ITAG5 and DPP4/CD26, whereas CD47^Lo^ cells did not.

**Table S1.** Summary of the participants in the study. There was no difference in the ages between the normal and OA group (*p*=0.13).

|  | **Total Number (Male and Female)** | **Age Range** |
| --- | --- | --- |
| **Normal** | **18 (14M/4F)** | **19-77** |
| **OA** | **15 (5M/10F)** | **46-83** |

**Table S2.** Summary of all clonal lines derived from all patients included in the study.

**Grey highlight:** Patient sample did not yield any cell growth following single cell sorting.

**Blue highlight:** Clonal cell line yielded enough number of cells for full characterization.

| **Patient Type** | **Age** | **Sex** | **Number of clones established** | **Cell line name** | **Final cell count** | **Population doublings** |
| --- | --- | --- | --- | --- | --- | --- |
| **Synovial Biopsies from Normal Individuals** | | | | | | |
| **Normal** | **77** | **M** | **3** | **A:C8** | **<30,000** | **<14.86** |
|  |  |  |  | **B:G1** | **<30,000** | **<14.86** |
|  |  |  |  | **A:B11** | **<30,000** | **<14.86** |
| **Normal** | **67** | **M** | **2** | **B:F6** | **<30,000** | **<14.86** |
|  |  |  |  | **B:G2** | **<30,000** | **<14.86** |
| **Normal** | **27** | **M** | **45** | **B:E8** | **121,000** | **16.87** |
|  |  |  |  | **B:D4** | **5,819,000** | **22.46** |
|  |  |  |  | **A:B10** | **945,000** | **19.84** |
| **Normal** | **77** | **M** | **3** | **A:C2** | **<30,000** | **<14.86** |
|  |  |  |  | **B:A2** | **<30,000** | **<14.86** |
|  |  |  |  | **B:A11** | **<30,000** | **<14.86** |
| **Normal** | **37** | **M** | **0** | **-** | **-** | **-** |
| **Normal** | **54** | **F** | **0** | **-** | **-** | **-** |
| **Normal** | **37** | **M** | **1** | **B:G10** | **<150** | **7.22** |
| **Normal** | **72** | **M** | **0** | **-** | **-** | **-** |
| **Normal** | **76** | **M** | **0** | **-** | **-** | **-** |
| **Normal** | **60** | **F** | **0** | **-** | **-** | **-** |
| **Normal** | **77** | **M** | **9** | **A:D12** | **1,767,000** | **20.74** |
|  |  |  |  | **A:C9** | **1,063,000** | **20** |
|  |  |  |  | **A:B3** | **937,000** | **19.83** |
|  |  |  |  | **A:B11** | **1,220,000** | **20.20** |
|  |  |  |  | **B:B7** | **1,397,000** | **20.40** |
|  |  |  |  | **B:B6** | **654,000** | **19.31** |
|  |  |  |  | **B:E9** | **1,116,000** | **20.1** |
|  |  |  |  | **A:G2** | **49,800** | **15.6** |
|  |  |  |  | **A:B12** | **406,000** | **18.34** |
| **Normal** | **62** | **M** | 16 | **B:F12** | **83,600** | **16.34** |
|  |  |  |  | **B:C11** | **335,200** | **18.34** |
|  |  |  |  | **A:G2** | **49,800** | **15.6** |
|  |  |  |  | **B:B4** | **335,200** | **18.34** |
|  |  |  |  | **B:G3** | **<150** | **7.22** |
|  |  |  |  | **B:F3** | **83,500** | **16.33** |
|  |  |  |  | **B:F7** | **69,600** | **16.1** |
|  |  |  |  | **A:D12** | **2,026,000** | **20.94** |
|  |  |  |  | **A:C11** | **<150** | **7.22** |
|  |  |  |  | **A:B10** | **<150** | **7.22** |
|  |  |  |  | **A:A7** | **33,200** | **15.01** |
|  |  |  |  | **A:C4** | **23,400** | **17.82** |
|  |  |  |  | **A:E3** | **<150** | **7.22** |
|  |  |  |  | **A:C1** | **<150** | **7.22** |
|  |  |  |  | **A:B1** | **<150** | **7.22** |
|  |  |  |  | **A:B3** | **<150** | **7.22** |
| **Normal** | **46** | **F** | **4** | **B:D3** | **12,800** | **13.64** |
|  |  |  |  | **B:A7** | **14,100** | **17.1** |
|  |  |  |  | **A:C9** | **1,601,000** | **20.59** |
|  |  |  |  | **A:E1** | **2,106,000** | **20.99** |
| **Normal** | **67** | **M** | **0** | **-** | **-** | **-** |
| **Normal** | **49** | **M** | **0** | **-** | **-** | **-** |
| **Normal** | **70** | **F** | **0** | **-** | **-** | **-** |
| **Normal** | **19** | **M** | **22** | **B:A2** | **2,778,000** | **21.4** |
|  |  |  |  | **B:B9** | **2,300,000** | **21.12** |
|  |  |  |  | **B:B10** | **4,174,000** | **21.98** |
|  |  |  |  | **B:E2** | **1,636,000** | **20.63** |
|  |  |  |  | **B:F3** | **1,832,000** | **20.8** |
|  |  |  |  | **B:G1** | **3,279,000** | **21.63** |
|  |  |  |  | **B:F12** | **1,195,000** | **20.17** |
|  |  |  |  | **B:F9** | **<50** | **5.64** |
|  |  |  |  | **B:G5** | **1,222,000** | **20.20** |
|  |  |  |  | **B:G12** | **1,668,000** | **20.66** |
|  |  |  |  | **B:F1** | **2,216,000** | **21.1** |
|  |  |  |  | **B:E10** | **1,886,000** | **20.83** |
|  |  |  |  | **B:B11** | **<50** | **5.64** |
|  |  |  |  | **B:E4** | **1,845,000** | **20.80** |
|  |  |  |  | **B:C12** | **1,888,000** | **20.83** |
|  |  |  |  | **B:C8** | **1,211,000** | **20.2** |
|  |  |  |  | **C:A10** | **335,200** | **18.34** |
|  |  |  |  | **C:B7** | **33,200** | **15.01** |
|  |  |  |  | **A:A7** | **1,295,000** | **20.3** |
|  |  |  |  | **A:C3** | **2,462,000** | **21.21** |
|  |  |  |  | **A:D3** | **335,200** | **18.34** |
|  |  |  |  | **A:F11** | **1,275,000** | **20.70** |
| **Normal** | **76** | **M** | **13** | **A:12** | **<50** | **5.64** |
|  |  |  |  | **B:H5** | **60,300** | **15.87** |
|  |  |  |  | **B:G9** | **33,200** | **15.01** |
|  |  |  |  | **B:D7** | **<50** | **5.64** |
|  |  |  |  | **B:C11** | **<50** | **5.64** |
|  |  |  |  | **A:G3** | **335,200** | **18.34** |
|  |  |  |  | **A:G4** | **4490** | **12.12** |
|  |  |  |  | **A:G10** | **<50** | **5.64** |
|  |  |  |  | **A:F6** | **996,000** | **19.91** |
|  |  |  |  | **A:F9** | **<50** | **5.64** |
|  |  |  |  | **A:E4** | **33,200** | **15.01** |
|  |  |  |  | **A:E8** | **239,000** | **17.85** |
|  |  |  |  | **A:C10** | **335,200** | **18.34** |
|  |  |  |  | **A:F12** | **40,600** | **15.30** |
| **Synovial Biopsies from OA Patients** | | | | | | |
| **OA** | **83** | **F** | **21** | **A:A7** | **75** | **6.22** |
|  |  |  |  | **A:E5** | **108** | **6.75** |
|  |  |  |  | **A:F7** | **325** | **8.34** |
|  |  |  |  | **A:F9** | **350** | **8.45** |
|  |  |  |  | **A:G8** | **675** | **9.39** |
|  |  |  |  | **B:A3** | **<50** | **5.46** |
|  |  |  |  | **B:A12** | **<50** | **5.46** |
|  |  |  |  | **B:C3** | **175** | **7.44** |
|  |  |  |  | **B:E5** | **150** | **7.22** |
|  |  |  |  | **B:F1** | **<50** | **5.46** |
|  |  |  |  | **B:G9** | **140** | **7.12** |
|  |  |  |  | **B:H4** | **1,442,100** | **20.45** |
|  |  |  |  | **A:D4** | **<50** | **5.46** |
|  |  |  |  | **A:C4** | **100** | **6.64** |
|  |  |  |  | **A:B10** | **<50** | **5.46** |
|  |  |  |  | **A:F1** | **75** | **6.22** |
|  |  |  |  | **A:F10** | **<50** | **5.46** |
|  |  |  |  | **A:G12** | **150** | **7.22** |
|  |  |  |  | **A:H11** | **<50** | **5.46** |
|  |  |  |  | **A:H12** | **<50** | **5.46** |
|  |  |  |  | **A:H3** | **46,900** | **15.59** |
| **OA** | **46** | **F** | **12** | **B:B10** | **597,900** | **19.18** |
|  |  |  |  | **B:B4** | **1,983,800** | **20.91** |
|  |  |  |  | **B:B2** | **968,700** | **19.87** |
|  |  |  |  | **B:E4** | **865,300** | **19.71** |
|  |  |  |  | **B:C9** | **904,000** | **19.77** |
|  |  |  |  | **B:D7** | **3,117,000** | **21.56** |
|  |  |  |  | **B:A7** | **1,814,300** | **20.78** |
|  |  |  |  | **B:E7** | **2,170,000** | **21.03** |
|  |  |  |  | **B:B6** | **1,697,960** | **20.68** |
|  |  |  |  | **B:A3** | **1,204,000** | **20.19** |
|  |  |  |  | **B:C10** | **3,236,200** | **21.61** |
|  |  |  |  | **B:D5** | **2,259,000** | **21.1** |
|  |  |  |  | **B:F12** | **12,800** | **13.64** |
| **OA** | **76** | **M** | **0** | **-** | **-** | **-** |
| **OA** | **66** | **M** | **0** | **-** | **-** | **-** |
| **OA** | **77** | **F** | **0** | **-** | **-** | **-** |
| **OA** | **66** | **F** | **17** | **B:H7** | **3,570,000** | **21.75** |
|  |  |  |  | **B:E4** | **1,094,000** | **20.04** |
|  |  |  |  | **B:C6** | **2,583,000** | **21.28** |
|  |  |  |  | **B:C10** | **1,506,000** | **20.51** |
|  |  |  |  | **B:D2** | **2,007,000** | **20.92** |
|  |  |  |  | **B:E7** | **150** | **7.22** |
|  |  |  |  | **B:C2** | **150** | **7.22** |
|  |  |  |  | **A:C2** | **1,881,000** | **20.83** |
|  |  |  |  | **A:B1** | **150** | **7.22** |
|  |  |  |  | **A:B3** | **2,903,000** | **21.46** |
|  |  |  |  | **A:E4** | **150** | **7.22** |
|  |  |  |  | **A:F5** | **2,763,000** | **21.39** |
|  |  |  |  | **A:C7** | **2,014,000** | **20.93** |
|  |  |  |  | **A:G9** | **2,067,000** | **20.97** |
|  |  |  |  | **A:F9** | **3,492,000** | **21.72** |
|  |  |  |  | **A:B3** | **2,713,000** | **21.36** |
|  |  |  |  | **A:A8** | **2,606,000** | **21.30** |
|  |  |  |  | **A:G8** | **150** | **7.22** |
| **OA** | **52** | **F** | **7** | **B:C1** | **33,200** | **15.01** |
|  |  |  |  | **B:C2** | **1,191,800** | **20.17** |
|  |  |  |  | **B:H7** | **1,191,800** | **20.17** |
|  |  |  |  | **B:B9** | **335,200** | **18.34** |
|  |  |  |  | **B:F12** | **63,300** | **15.94** |
|  |  |  |  | **B:E8** | **150** | **7.22** |
|  |  |  |  | **A:C3** | **150** | **7.22** |
| **OA** | **58** | **M** | **9** | **A:A7** | **150** | **7.22** |
|  |  |  |  | **A:B6** | **1,147,000** | **20.12** |
|  |  |  |  | **A:C9** | **627,500** | **19.25** |
|  |  |  |  | **A:D1** | **5,980** | **12.54** |
|  |  |  |  | **A:H3** | **7,000** | **12.76** |
|  |  |  |  | **A:H11** | **33,200** | **15.01** |
|  |  |  |  | **B:E1** | **1,812,000** | **20.78** |
|  |  |  |  | **B:G5** | **87,500** | **16.41** |
|  |  |  |  | **B:G12** | **39,900** | **15.28** |
| **OA** | **72** | **F** | **11** | **A:B7** | **150** | **7.22** |
|  |  |  |  | **A:C3** | **150** | **7.22** |
|  |  |  |  | **A:E6** | **34,500** | **15.1** |
|  |  |  |  | **B:C1** | **150** | **7.22** |
|  |  |  |  | **B:B4** | **13,600** | **13.72** |
|  |  |  |  | **B:A6** | **150** | **7.22** |
|  |  |  |  | **B:A7** | **1,456,000** | **20.46** |
|  |  |  |  | **B:A9** | **109,000** | **16.72** |
|  |  |  |  | **B:B11** | **817,500** | **19.62** |
|  |  |  |  | **B:C3** | **1,301,000** | **20.3** |
|  |  |  |  | **B:C5** | **1,125,000** | **20.1** |
| **OA** | **65** | **M** | **4** | **A:F8** | **15,200** | **13.9** |
|  |  |  |  | **A:B8** | **1,260,000** | **20.25** |
|  |  |  |  | **A:B6** | **150** | **7.22** |
|  |  |  |  | **A:D4** | **1,000,400** | **19.92** |
| **OA** | **72** | **F** | **0** | **-** | **-** | **-** |
| **OA** | **76** | **M** | **0** | **-** | **-** | **-** |
| **OA** | **75** | **F** | **0** | **-** | **-** | **-** |
| **OA** | **56** | **F** | **0** | **-** | **-** | **-** |
| **OA** | **61** | **F** | **11** | **D8** | **10,800** | **13.4** |
|  |  |  |  | **B5** | **150** | **7.22** |
|  |  |  |  | **B:C1** | **33,900** | **15.04** |
|  |  |  |  | **B:C5** | **332,000** | **18.33** |
|  |  |  |  | **B:D5** | **4,250** | **12.04** |
|  |  |  |  | **A:C12** | **27,900** | **11.43** |
|  |  |  |  | **A:H10** | **2,479,000** | **21.23** |
|  |  |  |  | **A:B7** | **12,100** | **13.55** |
|  |  |  |  | **A:B1** | **1,053,000** | **19.99** |
|  |  |  |  | **B:B5** | **33,100** | **15** |
|  |  |  |  | **B:B1** | **4,185,000** | **21.98** |

**Table S3.** Summary of the self-renewal capacity (population doublings) from all clonal lines derived in the study.

| **Population Doublings** | **<10** | **10 - 15** | **15 - 18** | **18 - 20** | **>20** |
| --- | --- | --- | --- | --- | --- |
| **# Normal Clones** | **69** | **54** | **45** | **31** | **29** |
| **# OA Clones** | **94** | **53** | **39** | **36** | **37** |

**Table S4.** Summary of the cell surface marker expression (in situ and in vitro) and differentiation potential of clones derived from normal individuals.

| ***Cell Potency*** | **Cell Surface Marker Expression In-Situ** | | | | | **Chondrogenic Capacity** | **Osteogenic Capacity** | **Adipogenic Capacity** | **Cell Surface Marker Expression In-Vitro** | | | | |
| --- | --- | --- | --- | --- | --- | --- | --- | --- | --- | --- | --- | --- | --- |
|  | **CD90** | **CD44** | **CD73** | **CD105** | **CD271** |  |  |  | **CD90** | **CD44** | **CD73** | **CD105** | **CD271** |
| MPCs (3) | Negative | Positive | Positive | Negative | Negative | Positive | Positive | Positive | Positive | Positive | Positive | Positive | Negative |
|  | Positive | Positive | Positive | Negative | Negative | Positive | Positive | Positive | Positive | Positive | Positive | Positive | Negative |
|  | Positive | Positive | Positive | Negative | Negative | Positive | Positive | Positive | Positive | Positive | Positive | Positive | Negative |
| Bi- potent Progenitors (5) | Positive | Positive | Positive | Negative | Negative | Negative | Positive | Positive | Positive | Positive | Positive | Positive | Negative |
|  | Positive | Positive | Positive | Negative | Negative | Positive | Negative | Positive | Positive | Positive | Positive | Positive | Negative |
|  | Negative | Positive | Positive | Negative | Negative | Positive | Negative | Positive | Positive | Positive | Positive | Positive | Negative |
|  | Negative | Positive | Positive | Negative | Negative | Positive | Negative | Positive | Positive | Positive | Positive | Positive | Negative |
|  | Negative | Positive | Positive | Negative | Negative | Positive | Positive | Negative | Positive | Positive | Positive | Positive | Negative |
| Uni- potent Progenitors (6) | Negative | Positive | Positive | Negative | Negative | Positive | Negative | Negative | Positive | Positive | Positive | Positive | Negative |
|  | Negative | Positive | Positive | Negative | Negative | Negative | Negative | Positive | Positive | Positive | Positive | Positive | Negative |
|  | Negative | Positive | Positive | Negative | Negative | Negative | Negative | Positive | Positive | Positive | Positive | Positive | Negative |
|  | Negative | Positive | Positive | Negative | Negative | Negative | Negative | Positive | Positive | Positive | Positive | Positive | Negative |
|  | Negative | Positive | Positive | Negative | Negative | Negative | Positive | Negative | Positive | Positive | Positive | Positive | Negative |
|  | Positive | Positive | Positive | Negative | Negative | Negative | Negative | Positive | Positive | Positive | Positive | Positive | Negative |
| No differentiation capacity (4) | Positive | Positive | Positive | Negative | Negative | Negative | Negative | Negative | Positive | Positive | Positive | Positive | Negative |
|  | Negative | Positive | Positive | Negative | Negative | Negative | Negative | Negative | Positive | Positive | Positive | Positive | Negative |
|  | Negative | Positive | Positive | Negative | Negative | Negative | Negative | Negative | Positive | Positive | Positive | Positive | Negative |
|  | Negative | Positive | Negative | Negative | Negative | Negative | Negative | Negative | Positive | Positive | Positive | Positive | Negative |

**Table S5.** Summary of the cell surface marker expression (in situ and in vitro) and differentiation potential of clones derived from OA patients.

| ***Cell Potency*** | **Cell Surface Marker Expression In-Situ** | | | | | **Chondrogenic Capacity** | **Osteogenic Capacity** | **Adipogenic Capacity** | **Cell Surface Marker Expression In-Vitro** | | | | |
| --- | --- | --- | --- | --- | --- | --- | --- | --- | --- | --- | --- | --- | --- |
|  | **CD90** | **CD44** | **CD73** | **CD105** | **CD271** |  |  |  | **CD90** | **CD44** | **CD73** | **CD105** | **CD271** |
| MPCs (13) | Positive | Positive | Positive | Negative | Negative | Positive | Positive | Positive | Positive | Positive | Positive | Positive | Negative |
|  | Positive | Positive | Positive | Negative | Negative | Positive | Positive | Positive | Positive | Positive | Positive | Positive | Negative |
|  | Positive | Positive | Positive | Negative | Negative | Positive | Positive | Positive | Positive | Positive | Positive | Positive | Negative |
|  | Positive | Positive | Positive | Negative | Negative | Positive | Positive | Positive | Positive | Positive | Positive | Positive | Negative |
|  | Positive | Positive | Positive | Negative | Negative | Positive | Positive | Positive | Positive | Positive | Positive | Positive | Negative |
|  | Positive | Positive | Positive | Negative | Negative | Positive | Positive | Positive | Positive | Positive | Positive | Positive | Negative |
|  | Negative | Positive | Positive | Negative | Negative | Positive | Positive | Positive | Positive | Positive | Positive | Positive | Negative |
|  | Negative | Positive | Positive | Negative | Negative | Positive | Positive | Positive | Positive | Positive | Positive | Positive | Negative |
|  | Negative | Positive | Positive | Negative | Negative | Positive | Positive | Positive | Positive | Positive | Positive | Positive | Negative |
|  | Negative | Positive | Positive | Negative | Negative | Positive | Positive | Positive | Positive | Positive | Positive | Positive | Negative |
|  | Negative | Positive | Positive | Negative | Negative | Positive | Positive | Positive | Positive | Positive | Positive | Positive | Negative |
|  | Negative | Positive | Negative | Negative | Negative | Positive | Positive | Positive | Positive | Positive | Positive | Positive | Negative |
|  | Negative | Positive | Positive | Positive | Positive | Positive | Positive | Positive | Positive | Positive | Positive | Positive | Positive |
| Bi- potent Progenitors (9) | Positive | Positive | Positive | Negative | Negative | Positive | Negative | Positive | Positive | Positive | Positive | Positive | Negative |
|  | Positive | Positive | Positive | Negative | Negative | Positive | Negative | Positive | Positive | Positive | Positive | Positive | Negative |
|  | Positive | Positive | Positive | Negative | Negative | Positive | Negative | Positive | Positive | Positive | Positive | Positive | Negative |
|  | Positive | Positive | Positive | Positive | Negative | Positive | Negative | Positive | Positive | Positive | Positive | Positive | Negative |
|  | Negative | Positive | Positive | Negative | Negative | Positive | Negative | Positive | Positive | Positive | Positive | Positive | Negative |
|  | Negative | Positive | Negative | Negative | Negative | Positive | Positive | Negative | Positive | Positive | Positive | Positive | Negative |
|  | Positive | Positive | Positive | Positive | Negative | Positive | Positive | Negative | Positive | Positive | Positive | Positive | Negative |
|  | Positive | Positive | Positive | Negative | Negative | Negative | Positive | Positive | Positive | Positive | Positive | Positive | Negative |
|  | Negative | Positive | Negative | Negative | Negative | Negative | Positive | Positive | Positive | Positive | Positive | Positive | Negative |
| Uni- potent Progenitors (9) | Positive | Negative | Positive | Negative | Negative | Positive | Negative | Negative | Positive | Positive | Positive | Positive | Negative |
|  | Positive | Positive | Positive | Negative | Negative | Negative | Positive | Negative | Positive | Positive | Positive | Positive | Negative |
|  | Negative | Positive | Positive | Negative | Negative | Negative | Negative | Positive | Positive | Positive | Positive | Positive | Negative |
|  | Positive | Positive | Positive | Negative | Negative | Negative | Negative | Positive | Positive | Positive | Positive | Positive | Negative |
|  | Positive | Positive | Positive | Negative | Negative | Negative | Negative | Positive | Positive | Positive | Positive | Positive | Negative |
|  | Positive | Positive | Positive | Negative | Negative | Negative | Positive | Negative | Positive | Positive | Positive | Positive | Negative |
|  | Positive | Positive | Positive | Negative | Negative | Positive | Negative | Negative | Positive | Positive | Positive | Positive | Negative |
|  | Positive | Positive | Positive | Positive | Negative | Negative | Negative | Positive | Positive | Positive | Positive | Positive | Negative |
|  | Positive | Positive | Positive | Positive | Negative | Negative | Negative | Positive | Positive | Positive | Positive | Positive | Negative |
|  | Negative | Positive | Positive | Negative | Negative | Negative | Negative | Positive | Positive | Positive | Positive | Positive | Negative |
| No differentiation capacity (2) | Negative | Positive | Positive | Negative | Negative | Negative | Negative | Negative | Positive | Positive | Positive | Positive | Negative |
|  | Positive | Positive | Positive | Negative | Negative | Negative | Negative | Negative | Positive | Positive | Positive | Positive | Negative |

**Table S6.** Summary of the cell surface marker expression (in situ and in vitro) and differentiation potential of clones derived from normal individuals.

| ***Cell Potency*** | **Patient** | **CD90^+^CD44^+^CD73^+^** | **Chondrogenic Capacity** | **Osteogenic Capacity** | **Adipogenic Capacity** | **Cell Surface Marker Expression In-Vitro** | | | | |
| --- | --- | --- | --- | --- | --- | --- | --- | --- | --- | --- |
|  |  |  |  |  |  | **CD90** | **CD44** | **CD73** | **CD105** | **CD271** |
| **Normal Synovium** | **1** | **Yes** | **Negative** | **Positive** | **Positive** | **Positive** | **Positive** | **Positive** | **Positive** | **Negative** |
|  | **1** | **No** | **Negative** | **Negative** | **Negative** | **Positive** | **Positive** | **Positive** | **Positive** | **Negative** |
|  | **2** | **Yes** | **Positive** | **Negative** | **Positive** | **Positive** | **Positive** | **Positive** | **Positive** | **Negative** |
|  | **2** | **No** | **Negative** | **Negative** | **Positive** | **Positive** | **Positive** | **Positive** | **Positive** | **Negative** |
|  | **3** | **Yes** | **Positive** | **Positive** | **Positive** | **Positive** | **Positive** | **Positive** | **Positive** | **Negative** |
|  | **3** | **No** | **Positive** | **Negative** | **Negative** | **Positive** | **Positive** | **Positive** | **Positive** | **Negative** |
|  | **4** | **Yes** | **Negative** | **Positive** | **Positive** | **Positive** | **Positive** | **Positive** | **Positive** | **Negative** |
|  | **4** | **No** | **Positive** | **Negative** | **Positive** | **Positive** | **Positive** | **Positive** | **Positive** | **Negative** |
| **OA Synovium** | **1** | **Yes** | **Negative** | **Positive** | **Positive** | **Positive** | **Positive** | **Positive** | **Positive** | **Negative** |
|  | **1** | **No** | **Negative** | **Negative** | **Positive** | **Positive** | **Positive** | **Positive** | **Positive** | **Negative** |
|  | **2** | **Yes** | **Positive** | **Negative** | **Negative** | **Positive** | **Positive** | **Positive** | **Positive** | **Negative** |
|  | **2** | **No** | **Positive** | **Positive** | **Positive** | **Positive** | **Positive** | **Positive** | **Positive** | **Negative** |
|  | **3** | **Yes** | **Negative** | **Negative** | **Positive** | **Positive** | **Positive** | **Positive** | **Positive** | **Negative** |
|  | **3** | **No** | **Positive** | **Positive** | **Positive** | **Positive** | **Positive** | **Positive** | **Positive** | **Negative** |
|  | **4** | **Yes** | **Negative** | **Positive** | **Positive** | **Positive** | **Positive** | **Positive** | **Positive** | **Negative** |
|  | **4** | **No** | **Negative** | **Negative** | **Negative** | **Positive** | **Positive** | **Positive** | **Positive** | **Negative** |
